# Real-time noise cancellation for EEG monitoring and brain–computer interfaces

**DOI:** 10.64898/2026.10.16.759581

**Authors:** Matilda Makkonen, Olli-Pekka Kahilakoski, Johanna Metsomaa, Pantelis Lioumis, Risto J. Ilmoniemi, Tuomas P. Mutanen

**Affiliations:** Department of Neuroscience and Biomedical Engineering, Aalto University School of Science, Espoo, Finland; BioMag Laboratory, HUS Medical Imaging Center, Aalto University, University of Helsinki and Helsinki University Hospital, Helsinki, Finland

**Author notes:** Corresponding author: Matilda Makkonen.

## Abstract

Raw electroencephalography (EEG) data suffer from a low signal-to-noise ratio (SNR). Effective real-time cleaning of EEG would improve the robustness of EEG monitoring, brain–computer interfaces, and brain-state-dependent stimulation applications in both laboratory and real-life environments.

Here, we present a real-time source-estimate-utilizing noise-discarding algorithm (rtSOUND) for EEG noise suppression. We adapted the established offline SOUND algorithm for real-time use and implemented it into an open-source EEG-processing software. Instead of using conventional temporal filtering, SOUND operates the data with a spatial filter. This makes it particularly suitable for real-time applications, since each streamed data sample can be cleaned separately in near real-time. Unlike temporal filtering methods, SOUND does not make assumptions about the frequency content of noise and brain signals but assumes that the noise topographies differ from those generated by brain activity.

With an EEG forward model, rtSOUND cross-validates the EEG time series and estimates the spatial distribution of noise, resulting in a data-cleaning linear spatial filter. Each streamed EEG sample is processed synchronously, and the filter is periodically updated in an asynchronous parallel process. The algorithm rapidly updates the spatial filter in response to changes in the data, ensuring that noise reduction is both immediate and accurately adapted to the current EEG state. rtSOUND was validated with transcranial magnetic stimulation (TMS)–EEG and resting-state EEG datasets measured from 25 healthy subjects.

SNR of both TMS-evoked potentials and resting-state data consistently improved across various metrics. rtSOUND particularly benefitted channels with poor SNR, while retaining the waveforms in high-quality channels. The algorithm was fast enough to process data streamed at 5 kHz. rtSOUND improves the SNR of EEG in real time, providing a robust method to improve signal quality for brain-state-monitoring applications.

## 1. Introduction

Electroencephalography (EEG) offers a direct measure of brain activity. When neurons communicate, they produce synchronous post-synaptic currents that reflect large-scale action potential activity. These currents generate electric fields strong enough to be measurable as scalp potentials. Since its introduction, EEG has established itself as a paramount method for recording brain activity in both research and healthcare. It is vital for several clinical applications, including epilepsy [1] and sleep monitoring [2], as well as continuous state monitoring in intensive care units (ICUs). Recently, the significance of continuous EEG recording in research applications has also grown, especially in areas such as brain–computer interfaces (BCI) [3] and EEG-guided brain stimulation techniques. In the future, the evolution of artificial-intelligence (AI) algorithms [4] is likely to speed up this development. However, both established BCI and AI solutions are prone to effects of noise [5], emphasizing the importance of recording high-quality raw EEG signals [6].

Raw EEG data have an inherently low signal-to-noise ratio (SNR), which is a major challenge in both clinical and research settings. EEG measurements are prone to environmental noise, and other medical and experimental devices can interfere with the measurement. High-quality EEG depends on good skin–electrode contacts [7]. During prolonged measurements, these connections tend to degrade due to the drying of electrode gel and shifts in the EEG cap relative to the scalp.

Raw EEG data are often displayed as continuous time courses. This common visualization shapes our understanding of EEG signals, leading us to reduce noise in the time domain by filtering out frequencies typically associated with noise. Yet, this conventional method has its drawbacks, particularly for real-time analysis. Time-domain filtering requires an adequate time window to determine the frequency content, with longer windows enhancing the accuracy of frequency response at the cost of decreased responsiveness to non-stationarities in the data [8]. Moreover, real-time time-domain filtering may cause phase shifts, introduce signal delays, and struggle to keep up with the variability of EEG signals, all of which can impair the analysis. Acausal filtering, which depends on future data for estimating the current signal, can improve estimates of signal values. However, in real-time acausal filtering, future data might need to be predicted with methods such as autoregressive modeling [9, 10]. These predictions decrease in accuracy the further they extend into the future, posing a significant challenge for real-time processing.

In contrast, if EEG data is reframed as a series of instantaneous ‘snapshots’, each representing the voltage pattern on the scalp at a specific moment, a new family of methods emerges, spatial filtering (Fig. 1). These techniques process each data sample independently, allowing for a swift update of the spatial filter if non-stationarities are detected. This adaptability offers a significant advantage over time-domain filtering, as it enables the spatial filter to be modified rapidly in response to changes in the data, ensuring that noise reduction is both immediate and more accurately attuned to the current state of the EEG. In addition, spatial filtering is often faster than time-domain filtering that employs conventional sample-by-sample convolution, due to the smaller number of required calculation operations.

**Fig 1.**
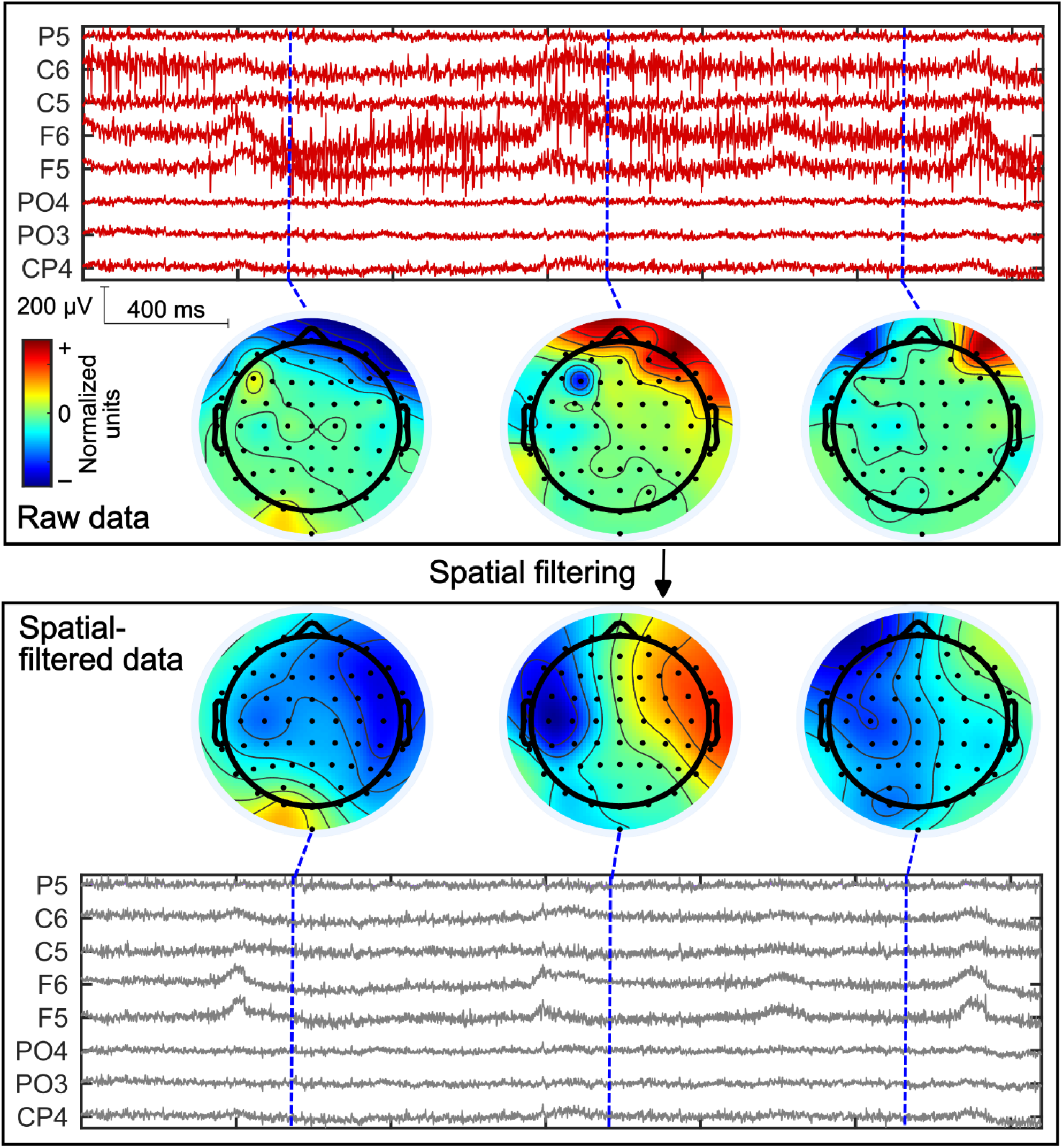
Schematic illustration of real-time spatial filtering of an EEG data stream. Upper box: Each time point of the raw EEG measurement (red curves) can be represented as a separate topography. The topographies illustrate the EEG voltage maps over the scalp. Lower box: When corrected with a noise-suppressing spatial filter, the topographies become smoother. Noise is assumed to be uncorrelated across EEG channels. If the time points corrected with the spatial filter are visualized as a continuous signal (grey curves), the noise in the signal time-series is also suppressed. For this visualization, the normalization of topographies was performed separately for raw and filtered data.

The key challenge is to develop a spatial filter that can isolate and eliminate noise-related spatial patterns or topographies, enabling efficient real-time noise suppression that could potentially overcome the constraints of temporal filtering methods. All useful filtering relies on *a priori* knowledge of the relevant signal characteristics and the nature of the remaining data, which may include noise, artifacts, or otherwise irrelevant signals. For example, low-pass filtering is appropriate when it is known that SNR is poor at high frequencies. It is important to recognize the assumptions made about the signal when removing noise or artifacts.

The source-estimate-utilizing noise-discarding algorithm (SOUND) [11, 12] is an established offline EEG cleaning method utilizing spatial filtering for noise suppression. The main assumption of SOUND is that due to the low-conductivity skull and the well-conducting skin, intracranially generated EEG signals are heavily correlated between neighbouring channels, whereas extracranially generated artifacts such as those from scalp muscles can be spatially less correlated. SOUND employs an EEG forward model to cross-validate the EEG-channel time points, estimating the spatial distribution of noise and resulting in a noise-suppressing linear spatial filter. SOUND assumes that noise topographies in EEG are spanned by a set of spatial patterns that is different from the set of topographies generated by neuronal activity.

In this work, we present a real-time version of the SOUND algorithm, rtSOUND, for EEG noise suppression. We conduct an extensive validation of the algorithm with both resting-state EEG [13-15] and transcranial magnetic stimulation-elicited EEG (TMS–EEG) [16]. TMS–EEG data were chosen for this study, because they are a prime example of noisy and artifactual data [16], revealing the possible limitations of preprocessing methods. On the other hand, finding effective methods to clean resting-state EEG in real time is essential for facilitating various EEG-guided applications, such as EEG-guided TMS [9, 10]. We demonstrate that rtSOUND improves SNR of both resting-state EEG and TMS−EEG data, streamed at the rate of thousands of samples per second, particularly benefitting the noisiest channels.

## 2. Materials and Methods

### 2.1. Real-time SOUND algorithm

As Fig. 1 illustrates, EEG data can be interpreted as a constant stream of topographies. Thus, the EEG signal can be written as a time-dependent NC ×1signal vector y(t), whose element *y*_i_(*t*)corresponds to the voltage measured by channel *i* at time *t, N*_*C*_ being the number of channels:

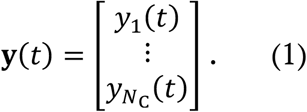

The measured signal ***y***is the sum of the voltage patterns resulting from active cortical current sources and noise:

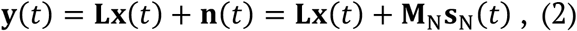

where ***L***is the NC×NS lead-field matrix [17] (_*NS*_ being the number of modelled cortical current sources), x(*t*) is an unknown *N*_S *×*_ 1 vector describing the active cortical current distribution at time *t*, and M_*N*_ and s_*N*_ are the *N*_*C ×*_ *N*_*N*_ noise mixing matrix and *N*_*N ×*_ 1 noise signal vector, respectively, defining the noise **n** at time *t*. In general, the number of noise components, *N*_*N*_, is unknown. M_*N*_ holds the scalp potential patterns of individual noise components in its columns, whereas the rows of s_*N*_ correspond to their time courses.

Our real-time noise cancellation method utilizes the SOUND algorithm [11, 12], which was first developed for offline cleaning of EEG [11]. Here, we use the SOUND algorithm to produce a spatial filter, W_SOUND_, to suppress the noise topography components (W_SOUND_M_N_ ≈ 0) while preserving the neuronal voltage patterns in each streaming EEG topography (W_SOU*N*D_L ≈ L). Here, we assume that compared to neuronal EEG, noise topographies are spanned by a different set of spatial components. Thus, we estimate the noise-free EEG signals ŷ(*t*) as

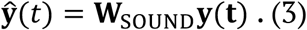

It turns out that if we know the noise covariance matrix, we can write the SOUND spatial filter as (see the detailed derivation in the Supplementary material)

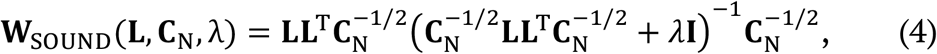

where c_*N*_ is the *N*_*C ×*_ *N*_*C*_ noise covariance matrix, and λ is the regularization parameter.

Eqs. 3 and 4 can be interpreted as a noise-suppressed projection of the EEG signal to the source space followed by a projection of the noise-suppressed neuronal signal estimates back to the original EEG sensor space. The noise-covariance terms in Eq. 4 work as a penalty in the EEG signal space, based on the structure of the noise; the signal-space directions in which the noise is more pronounced will be suppressed more. The regularization parameter λ defines how aggressively the noise is suppressed. A larger λ value means greater noise attenuation, but with the cost of possible suppression of neuronal signal components.

To use Eq. 4, we need to estimate the noise covariance from the streaming data. In the Supplementary material, we show that, assuming uncorrelated noise covariance across the channels, c_*N*_ can be estimated very efficiently from a data buffer [18]. Because we assume uncorrelated noise, the SOUND filter is particularly effective against isolated corrupted channels. However, previous studies have shown that SOUND suppresses noise effectively also if the noise topographies differ enough from the neuronal voltage patterns, even though the assumption of uncorrelated noise is not fully satisfied [11].

The workflow of rtSOUND is as follows (see Fig. 2 for a schematic). Before the measurement, the user provides a suitable lead-field matrix and sets the desired regularization value. When the measurement begins, each raw data sample is handled in a real-time (synchronous) process, including baseline correction and spatial filtering with the latest SOUND filter W_SOUND_. The SOUND filter is constantly updated in a parallel (asynchronous) process, based on the latest circular data buffer. The parallel update process ensures that rtSOUND reacts promptly to changes in noise distributions. For instance, if an EEG channel is suddenly disconnected, rtSOUND receives this information in the next noise-covariance estimation round, updating the SOUND filter accordingly.

**Fig 2.**
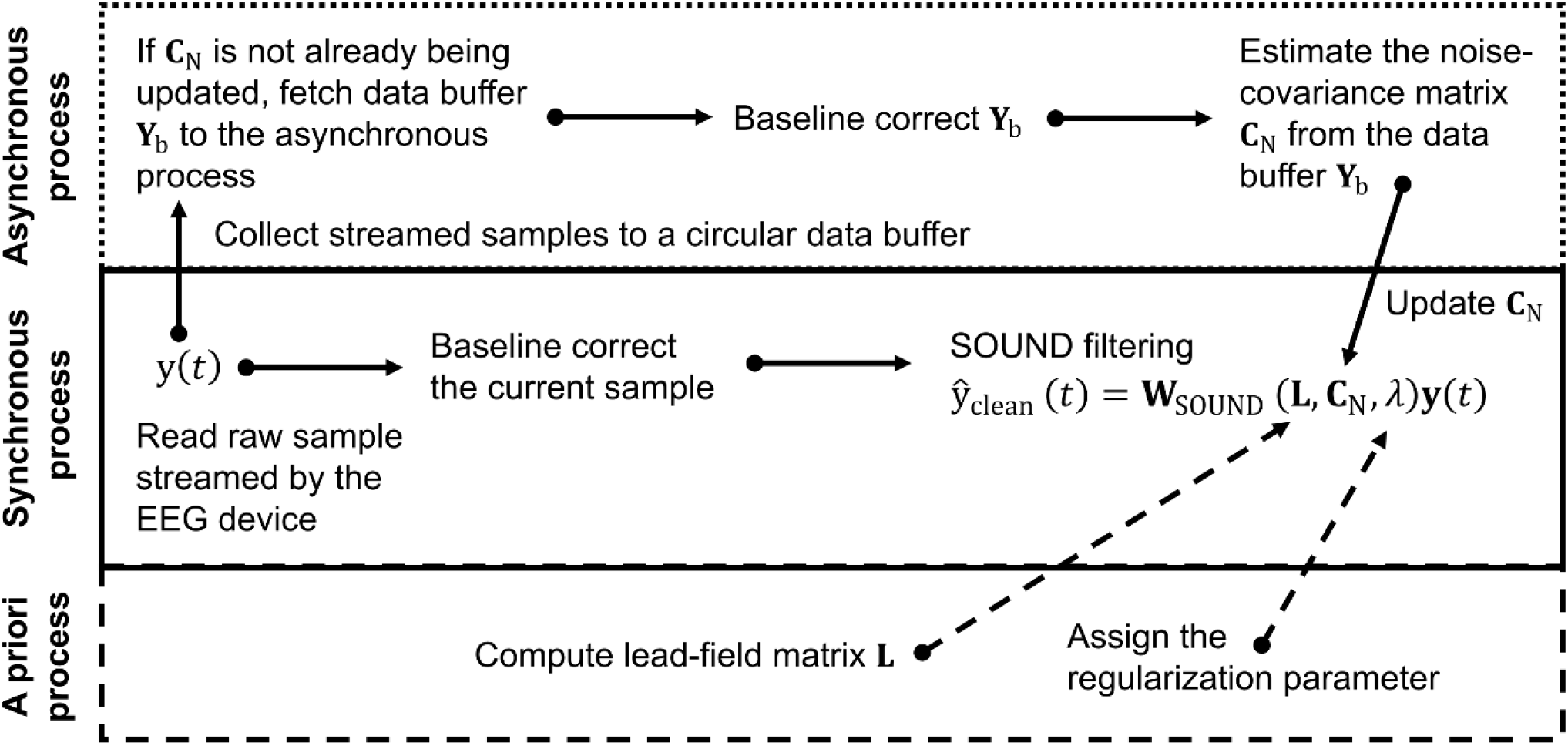
Schematic workflow of running rtSOUND. Before the measurement, the user provides a suitable lead-field matrix and a value of the regularization parameter λ. When the measurement begins, each sample of raw data is baseline corrected and multiplied with the latest rtSOUND filter W_SOUND_ in a real-time (synchronous) process. In a parallel (asynchronous) process, the rtSOUND filter is updated by estimating the new noise-covariance matrix c_*N*_ based on the baseline-corrected version of the latest data buffer.

It is important to note that, while rtSOUND requires a specialized parallel data-processing scheme (Fig. 2), all assumptions regarding offline SOUND also apply for rtSOUND. Thus, the major observations reported in earlier SOUND publications also hold for rtSOUND.

### 2.2. Software architecture for testing rtSOUND

rtSOUND was tested with the real-time EEG-processing software NeuroSimo, described in detail in [19]. The software is specifically designed for EEG-controlled TMS experiments, and it adopts a microservice architecture implemented using the Robot Operating System (ROS 2) [20]. In such an architecture, the software comprises multiple services that communicate with each other. Each service is responsible for a single, clearly defined task.

In NeuroSimo [19], one such service receives raw EEG data from the EEG device and stores it in a buffer. Another service, called *Preprocessor*, is provided access to this buffer for cleaning the streamed data samples. The data from *Preprocessor* is then transmitted to another service, *Decider*, which can execute algorithms such as real-time estimation of the phase of a specific neuronal oscillation. Based on these analyses, stimulation pulses can be delivered on a TMS device. With a multi-locus TMS device, the stimulation location can also be altered electronically or robotically [21, 22]. In the validation of rtSOUND with pre-recorded data, we employed NeuroSimo’s simulation mode, in which the data is entered into the data-processing pipeline, and the processing is performed as if the data were acquired in real time.

Although NeuroSimo is tailored for EEG-guided TMS, the present study primarily focuses on the performance of rtSOUND within *Preprocessor*. Preprocessing implemented within NeuroSimo allows integration with various applications supported by the software, including BCI.

An rtSOUND implementation is available online at https://github.com/matildamakkonen/rt-sound-neurosimo-plugin as a plug-in for the NeuroSimo software.

### 2.3. Constructing the lead-field matrix

The lead-field matrix was based on the open-source New York Head model [23], designed as a standard EEG forward model. The New York Head is derived from a nonlinear average of MRI scans from 152 adult human brains, known as the ICBM152 anatomical template. It includes six tissue types (scalp, skull, CSF, gray matter, white matter, and air cavities), segmented at 0.5-mm^3^ resolution. The model was solved for 231 standard electrode locations and 10016 cortically constrained dipolar current sources, oriented perpendicularly to the local cortical surface, using the finite-element method.

We identified the rows of the original New York Head lead-field matrix corresponding to our 10–20 electrode layout (see Section 2.4.1) and reordered them to match the input order of the EEG channels in the real-time data stream. To account for the effect of the reference channel on the measured neuronal topographies, we identified the channel in the New York Head closest to the actual reference electrode position (right mastoid) in the validation datasets and subtracted its lead field from each row of the pre-selected lead-field matrix.

Finally, we computed the left factorization of the singular-value decomposition of the lead-field matrix and used it in the rtSOUND algorithm, as suggested in [12]. This simplification to reduce the computational load of rtSOUND is valid because, strictly speaking, SOUND relies on the covariance of the lead fields rather than the original lead-field matrix [12], as suggested by Eq. 4.

### 2.4. Algorithm validation with pre-recorded data

#### 2.4.1. Pre-recorded data for validation

The algorithm was validated with a dataset consisting of resting-state EEG and TMS– EEG measured from 25 right-handed healthy individuals (11 females, 14 males, ages 28 ± 6.6 years). The dataset was collected as a part of the EU-funded ConnectToBrain project [24]. The study was accepted by the Coordinating Ethics Committee of the

Hospital District of Helsinki and Uusimaa and it followed the Declaration of Helsinki. During the experiment, the subjects sat on a comfortable chair and had their gaze fixated at a cross on the wall.

EEG was recorded at a sampling rate of 5000 Hz with a 24-bit biosignal amplifier (NeurOne, Bittium Plc, Oulu, Finland) in DC mode and a TMS-compatible 62-channel cap with Ag/AgCl sintered C-electrodes (EasyCap BC-TMS-64-X20-UCMW, EasyCap, Herrsching, Germany), placed according to the International 10–20 system. The reference electrode was on the right mastoid, and the ground electrode on the right cheekbone. Electrode impedances were kept below 5 kΩ, verified twice during the measurement.

For TMS, the navigated Nexstim NBS5 system (Nexstim Plc, Helsinki, Finland) with a figure-of-eight cooled coil (outer loop diameter 70 mm) delivered biphasic pulses consistently at the *abductor pollicis brevis* motor hotspot of the subject. Stimulation intensity was set at 110% of the resting motor threshold (RMT). RMT was defined as the intensity at which 10 out of 20 motor-evoked potentials had a peak-to-peak amplitude of least 50 μV. The average inter-stimulus interval was 4.25 s, with a random jitter of ± 0.25 s (uniform distribution). This setting was adjusted in the Nexstim system to random mode ranging between 4 and 4.5 seconds. The coil recharging artifact occurred at 1.9 s post-pulse.

T1-weighted MRI images of the participants were used for neuronavigation. Adaptive acoustic masking noise to the subjects was produced and controlled by the TAAC toolbox [25]. Before the main experiment, a quality check involving 20 pulses was performed to ensure the efficacy of noise masking, by visually confirming that no clear central positive response was observed in the time range 100−200 ms after the pulse [26]. The maximum permissible noise level was limited to 90 dB.

The measurement sequence started with two 5-minute resting-state EEG recordings. Then, the subject listened to the masking noise and the TMS coil was placed over the target location, but no pulses were given. This was followed by four 21-minute runs of TMS–EEG with 300 pulses each. Finally, another 5-minute resting-state EEG measurement was performed with the masking noise and the coil over the target.

From each participant, we utilized the first 5 minutes of EEG data from the fourth TMS–EEG measurement, as well as the entire last resting-state EEG measurement. These are later referred to the TMS–EEG data and resting-state data, respectively. These datasets were chosen since they were likely to correspond to the noisiest measurements.

#### 2.4.2. Real-time SOUND correction of pre-recorded validation data

The pre-recorded EEG data were formatted into comma-separated value (CSV) files and run through the NeuroSimo software described in Section 2.2. The software ran on a powerful laptop computer (Dell Precision 7680) with a 24-core Intel Core i9-13950HX processor, 32 GB of RAM and the Ubuntu 22.04 LTS operating system with the PREEMPT_RT kernel patch. The New York Head model-based lead field (see Section 2.3) was used for running rtSOUND for every participant’s data. The preprocessed data were saved by the software in CSV format.

The performance of rtSOUND depends on several tunable parameters that govern its responsiveness, stability, and signal preservation. Details on the parameters and their values used here are presented in the Supplementary material and further discussed in the Practical considerations subsection in the Discussion. Each dataset was run through the processing pipeline twice: on the first run, the SOUND filter calculation was performed normally, but the filter was not updated to the sample-by-sample matrix-multiplication process. This produced a dataset in which only the real-time baseline correction part of rtSOUND was applied. On the second run, the full rtSOUND was applied to the dataset. These two datasets were then compared to measure only the effect of rtSOUND while ruling out the possible effects of the baseline correction.

#### 2.4.3. Analysis of validation with pre-recorded data

To guide the validation of rtSOUND and evaluate its effectiveness, we defined four primary design aims for the algorithm: First, rtSOUND correction should improve the overall SNR. Second, since rtSOUND replaces signals in noisy channels with estimates based on all channels, these estimates must be accurate and reasonable. Third, the algorithm should rapidly respond to sporadic noise changes by updating the noise-suppressing spatial filter in real time. Finally, rtSOUND should focus noise suppression on low-SNR channels while leaving high-SNR channels largely unaltered to avoid overcorrection.

To evaluate whether these aims were achieved, we derived the following four research questions:

1. How does rtSOUND change SNR?
2. How similar is rtSOUND correction to gold-standard offline interpolation methods?
3. How fast and how robustly does rtSOUND correct sporadic noise signals?
4. How do the rtSOUND-elicited changes depend on the original SNR of the raw data channels, and to what extent does rtSOUND undesirably reduce the amplitudes of channels with high signal quality (*i*.*e*., low noise level)?

##### QUESTION 1: SNR CHANGE DUE TO RTSOUND

We calculated SNR for both the raw (real-time baseline-corrected) and rtSOUND-corrected signals in two ways. The first metric relied on channel-wise SNR estimation, based on filtered resting-state signal amplitudes and TMS-evoked potential (TEP) amplitudes. Continuous resting-state data were lowpass filtered at 40 Hz (representing the brain signal) and separately bandpass filtered between 55 and 95 Hz (representing the noise). These frequency bands were chosen to estimate the resulting SNR in the neuronally relevant frequencies (below 40 Hz), assuming that, apart from the 50-Hz line noise, the *a priori* estimate of the distribution of noise is approximately uniform across the frequencies. The channel-wise SNR was calculated as follows:

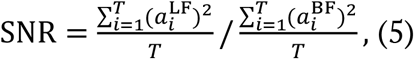

where *T* is the number of samples in the entire resting-state recording, and 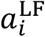 and 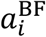 are the values of the *i*‘th sample of the lowpass- and bandpass-filtered signal *a*, respectively.

The TMS−EEG data were first epoched from –2 to +2 s around the TMS pulse, resulting in 57−70 trials per dataset, following the computation of TEPs as the average over epochs. The SNR metric for TEPs for each channel was defined as the ratio of the mean-square TEP (from 20 to 2000 ms after the TMS pulse) to the mean-square EEG signal from 2000 to 20 ms before the TMS pulse.

Our second metric utilized ICA and automatic component labelling [27] to estimate SNR. This metric was constructed to account also for those artifacts that overlapped with neuronal signals in the frequency domain. For calculating the ICA-based SNR, the resting-state dataset was baseline-corrected, re-referenced to global average and decomposed into 30 independent components with FastICA [28]. The TMS−EEG epochs were preprocessed by removing the baseline (2000 to 10 ms before the pulse), replacing the pulse artifact timespan (–2 to 10 ms) with zeros, and rereferencing the data to global average before running ICA for the concatenated epochs (from 20 to 2000 ms after the TMS pulse). The independent components were then classified automatically with the ICLabel toolbox [27]; those components that were classified as brain-originated with >90% probability were considered non-artifactual. All the remaining components were considered artifacts. SNR was defined as the ratio between the total variance of the non-artifactual components and the total variance of the artifactual components. To ensure that the selected probability threshold would not bias the results, the SNR values were also calculated for the TEP data with a threshold of 80%.

##### QUESTION 2: SIMILARITY OF RTSOUND AND INTERPOLATION

To assess the reliability of the rtSOUND predictions in the noisiest channels, we checked how similar rtSOUND correction is to gold-standard offline interpolation methods. We chose the three channels with the lowest SNR for each dataset based on the channel-wise SNR values. We divided the resting-state data into 1-s non-overlapping epochs. TMS−EEG data were epoched from 20 to 2000 ms after the TMS pulse. The chosen three channels were interpolated in each epoch with spherical-spline interpolation using the EEGLAB [29, 30] function eeg_interp with the default parameters. We calculated the time-correlation and relative root-mean-square deviation (RMSD) between the rtSOUND-corrected and interpolated signals. The relative RMSD was calculated as follows:

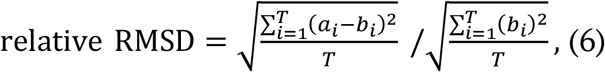

where *T* is the length of the signal in samples, and 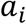 and 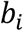 are the *i*’th samples of the interpolated and rtSOUND-corrected signals, respectively. We focused on the median and interquartile range of the time correlations and relative RMSD over epochs and over the three chosen channels. With the same metrics, we also compared the rtSOUND-corrected and raw signals.

##### QUESTION 3: CORRECTION OF SPORADIC NOISE

We investigated how fast and how robustly rtSOUND corrects sporadic noise signals by applying it to synthesized data. We created the simulated EEG data with a known noiseless ground truth, using the New York Head model, decimated to 10016 sources. We placed four neuronal sources (current dipoles) on the cortex, with oscillation frequencies drawn from Gaussian distributions centered at 8, 19, 30, and 41 Hz (1-Hz standard deviation) for the right primary visual cortex, left primary motor cortex, right supplementary motor area, and left prefrontal cortex, respectively. Each oscillator had a random phase relative to the onset of the simulation, and the maximum amplitude across EEG channels was set to 20 µV for each source. White noise was added, resulting in a power SNR of 10.

We then introduced sporadic line noise into the simulated ground-truth data. Each 50-Hz noise interval lasted 10 seconds and appeared at randomly chosen electrodes (1–4 at a time) at random times. The amplitude of each noisy channel was drawn from a Gaussian distribution centered at 50 µV (standard deviation 10 µV).

The noisy simulated data were then processed with rtSOUND, followed by manual annotation of the datasets with added noise. Based on visual inspection, we marked time points in which (1) a new noise signal was activated and (2) rtSOUND updated its correction in response to changes in the noise distribution (see Fig. 3 for a representative example).

**Fig 3.**
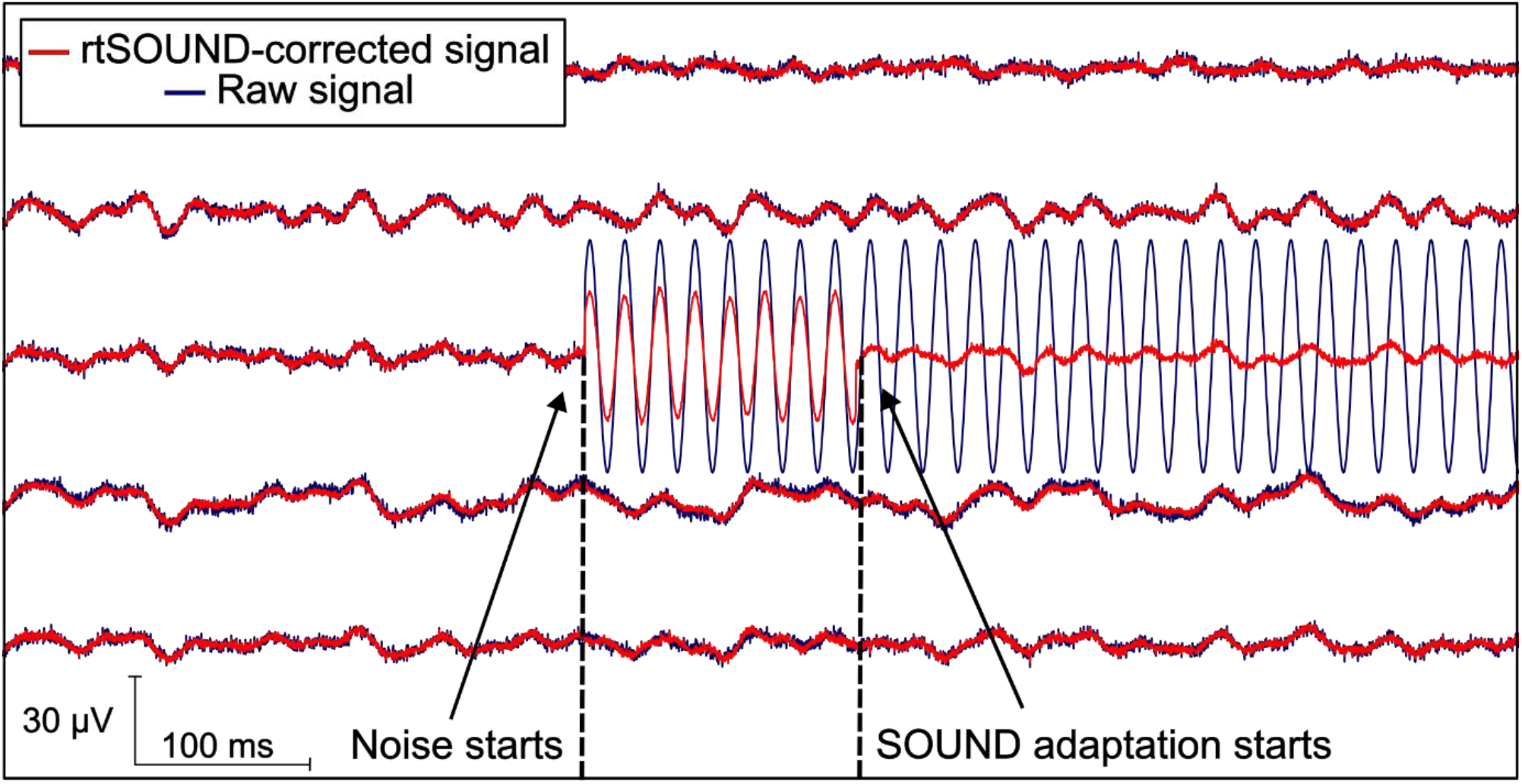
Example of manual annotation of data with added noise. The curves represent simulated data in five EEG channels. In the overlapping raw (blue) and rtSOUND-corrected (red) signals, we marked the time points in which the added noise started and rtSOUND correction adapted to the new noise distribution.

We calculated the relative RMSD for each noise instance (*N*=47) in the affected channel(s) for a 5-s segment after rtSOUND started correcting the noise. The RMSD was calculated both between the noisy and ground-truth (GT), and the rtSOUND-corrected and GT data. The difference in relative RMSD between the noisy−GT and rtSOUND−GT values was calculated for each noise instance.

##### QUESTION 4: RELATIONSHIP BETWEEN ORIGINAL SNR AND RTSOUND-ELICITED CHANGES

We built a linear mixed-effects (LME) model to detect a potential relationship between the initial SNR and the change in the signal amplitude after rtSOUND. The signal amplitude change for each channel was quantified as the relative mean-square deviation between the rtSOUND-corrected and raw signals (entire resting-state recording or the average TEP). In the LME model, the fixed linear effect of the initial SNR on the signal amplitude change was tested, while accounting for possible variability of the subjects and channels as random effects. Thus, the subject and EEG channel were modelled as random intercepts in the model: ChangeInSignal ∼ 1 +

InitialSNR + (1 | Subject) + (1 | Channel). We identified this as the maximal-random-effect structure for our experimental design [31], as only one value of SNR and change in signal exists for each subject−channel combination. The initial SNR and signal change values were log- and *z*-transformed before applying the LME model to ensure normal distribution of the residuals and to mitigate the influence of differing scales, facilitating model convergence and interpretation of model coefficients.

We also investigated how the initial channel-specific SNR affects the SOUND-induced SNR change. We analyzed this with Spearman correlation, because we assumed a monotonous, but not necessarily linear relationship between the variables. To test the amplitude suppression in the less noisy channels, we chose the 3 least noisy channels for each dataset based on the channel-specific SNR values. We divided the resting-state data into 1-s non-overlapping epochs, and epoched the TMS−EEG data from 20 to 2000 ms after the TMS pulse. We calculated the time correlations and relative RMSD between the rtSOUND-corrected and raw signals across epochs and the three chosen channels.

#### 2.5. Algorithm validation with real-time-streamed data

##### 2.5.1. Real-time-streamed validation data collection

The implementation of the rtSOUND algorithm was validated with real-time EEG measurements of two healthy right-handed subjects (25-year-old female and 35-year-old male; these subjects also belonged to the cohort for the pre-recorded validation data) who gave their written and informed consent. EEG was measured and sampled at 5000 Hz with a 24–bit biosignal amplifier in DC mode (NeurOne, Bittium, Oulu, Finland) and a TMS-compatible 62-channel cap with Ag/AgCl sintered C-electrodes (EasyCap BC-TMS-64-X20-UCMW, EasyCap, Herrsching, Germany), placed according to the International 10–20 system. The reference electrode for the EEG was placed on the right mastoid and the ground electrode on the right cheekbone. Electrode impedances were kept below 5 kΩ.

For both subjects, four 5-minute measurements were conducted:

1. Resting state, eyes open
2. Resting state, eyes closed
3. Resting state, eyes open, and added noise
4. Resting state, eyes closed, and added noise

In measurements 3 and 4, noise was added by gently pressing a randomly chosen EEG electrode with a finger for approximately 10 seconds at a time, changing the electrode after each 10-second period. This procedure introduced visible 50-Hz line noise in the manipulated channels. During the first minute of the recording, no noise was added. During the second minute of the recording, noise was added to one electrode at a time. During the third, fourth, and fifth minutes of the recording, noise was added to two, three, and four electrodes at a time, respectively.

rtSOUND was run during these measurements with identical parameters as with the pre-recorded validation data, the only difference being that the EEG data were streamed directly from the EEG device to the data-processing pipeline. While the streamed data was corrected in real-time with rtSOUND and saved for post-hoc analysis, also the raw data was saved. After the experiments, these raw signals were run through the real-time baseline-correction pipeline in simulation mode. This enabled comparison between rtSOUND-corrected data and signals that were baseline-corrected in an identical way. Thus, any difference observed in the post-hoc analysis could be attributed to rtSOUND. The same laptop computer with the same NeuroSimo software version was employed in all validation experiments.

#### 2.5.2. Analysis of validation with real-time-streamed data

In the validation with real-time-streamed data, we concentrated on the following research questions:

1. How fast is the rtSOUND correction for one sample in real time and for the filter update in the parallel process?
2. How much does rtSOUND improve SNR of real-time resting-state measurements?
3. How fast and how robustly does rtSOUND correct sporadic noise?

To answer Question 1, we measured and saved the single-sample processing durations on a per-sample basis and the SOUND filter update duration in all updates in all measurements. For Question 2, we calculated the ICA−SNR for both the raw and rtSOUND-corrected resting-state measurements.

To answer Question 3, we manually annotated the datasets with added noise in the same way as the simulated data (see Fig. 3). For each instance of added noise in one channel, we took 1 s of rtSOUND-corrected signal before the start of the added noise and calculated its mean power spectrum between 0 and 80 Hz to obtain a baseline measure. We also calculated the mean power spectra for the rtSOUND-corrected and raw signals from 0 to 1 s after rtSOUND had adapted to the new noise distribution. The power spectra were computed using Welch’s method with Hamming-windowed sections of 1024 samples each (with 50% overlap), followed by averaging the periodograms of these segments using a 1024-point Fast Fourier Transform. The resulting three spectra were compared with relative RMSD between the rtSOUND-corrected pre-noise and post-noise spectra, and between the rtSOUND-corrected and raw post-noise spectra.

### 2.6. Adding real-time eye-blink artifact removal

A factor that often disturbs real-time analysis of the EEG, in addition to noise, is due to large eye-blink artifacts. A previous study demonstrated that it is possible to remove blink artifacts in real time using a spatial filter based on eye-blink topographies extracted with ICA [32]. This spatial filter can be combined with the SOUND spatial filter in sample-by-sample correction to remove both noise and blink artifacts. Here, we wanted to test this approach in an actual real-time setting.

From the pre-recorded validation dataset, we chose five subjects whose EEG had large eye-blink artifacts. To mimic a realistic processing pipeline, the data from the first resting-state measurement that were not used in the validation runs (see Section 2.4.1) were used to calibrate the ICA filter. The calibration data were high-pass filtered from 1 Hz and decomposed into 30 components with FastICA [28]. The independent-component topographies corresponding to the eye blinks and eye movement were manually chosen, which resulted in two artifactual components being removed from each subject. An *N*_*C ×*_ *N*_*C*_ blink-suppressing spatial filter W_blink_ was built from the chosen topographies as follows:

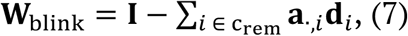

where 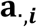 and 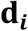 are the *i*‘th column and row of ICA mixing matrix A and demixing matrix D, respectively, and c_rem_ is the list of ICA component indices corresponding to the ocular-artifact components.

The blink-suppressing filter was introduced into the NeuroSimo *Preprocessor* pipeline. The script was modified so that each streamed sample was multiplied by the blink-removing filter in between the baseline correction and multiplication by the SOUND filter. The SOUND filter was calculated with the EEG data before blink correction. The resting-state and TMS−EEG datasets for each of the five subjects were then run through the pipeline.

The resulting data were analyzed with three metrics: the ICA−SNR metric, the durations of processing a single sample (aggregated over each second) and updating the SOUND filter, and a modified ICA-based eye-component metric. The latter was defined as the number of independent components that the automatic component classifier ICLabel [27] labelled as eye-artifact-components with >90% probability.

### 2.7. Analyzing the interaction between rtSOUND and TMS-evoked muscle artifacts

TMS-evoked muscle artifacts often mask the early TEP deflections, hindering TEP analysis. Muscle artifacts are somewhat correlated between EEG channels and thus do not entirely fulfill the assumptions of SOUND. However, as SOUND can also handle noise that does not fully satisfy the assumption of uncorrelatedness [11], we analyzed how SNR after and before rtSOUND is affected by large TMS-evoked muscle artifacts in the raw data.

We analyzed the 25 pre-recorded TMS−EEG datasets. Epochs were extracted from the only real-time baseline-corrected data (without rtSOUND) and preprocessed by removing the baseline (2000 to 10 ms before the pulse), interpolating the pulse artifact timespan (−2 to 10 ms) with cubic interpolation based on 1 ms of data before and after the interpolated time window, and re-referencing the data to global average before calculating the average TEP for each dataset. We calculated the global mean-field amplitude (GMFA) from each average TEP as the standard deviation over channels [33]. We quantified the muscle artifact as the maximum GMFA value in the range 11−50 ms after the pulse. The lower boundary 11 ms was set to avoid conducting the analysis with interpolated data.

The datasets were divided into two groups based on the muscle-artifact size. Datasets with a muscle-artifact amplitude of at least 50 μV in the GMFA were assigned to the large-muscle-artifact group, and the rest were assigned to the small-muscle-artifact group. The ICA−SNR values (see Section 2.4.3, Question 1) of the rtSOUND-corrected datasets in the two groups were then compared with the Wilcoxon rank-sum test [34]. This was repeated for the ICA−SNR values of only real-time baseline-corrected data to compare the difference between groups in both rtSOUND-corrected and raw data.

### 2.8. Quantifying the phase effect of rtSOUND

Many brain-state-dependent stimulation paradigms [10, 35] include real-time phase analysis of EEG oscillations for targeting stimulation. We investigated how rtSOUND affects the accuracy of phase estimation in simulated data, which allowed comparisons with a true ground-truth (GT) signal (see Section 2.4.3, Question 3 for details on the simulated data). For each added noise instance, we created 1-s epochs after rtSOUND started correcting the noise. This was performed for three datasets: simulated GT data, GT with added noise, and the rtSOUND-corrected noisy data. We investigated the effects on phase in the C3 channel and the 19-Hz frequency, since the simulated data included an oscillation centered at 19 Hz originating from the left primary motor cortex. We obtained the phase over each epoch with the Fast Fourier Transform and calculated the phase difference between GT and the noisy signal, and between GT and the rtSOUND-corrected signal. We then calculated the circular median and range of phase differences over epochs.

## 3. Results

### 3.1. Results of validation with pre-recorded data

Overall, rtSOUND detected and notably suppressed noise. Representative examples of the ability of rtSOUND to remove noise from the resting-state EEG signal and TEP are presented in Figs. 4 and 5, respectively.

**Fig 4.**
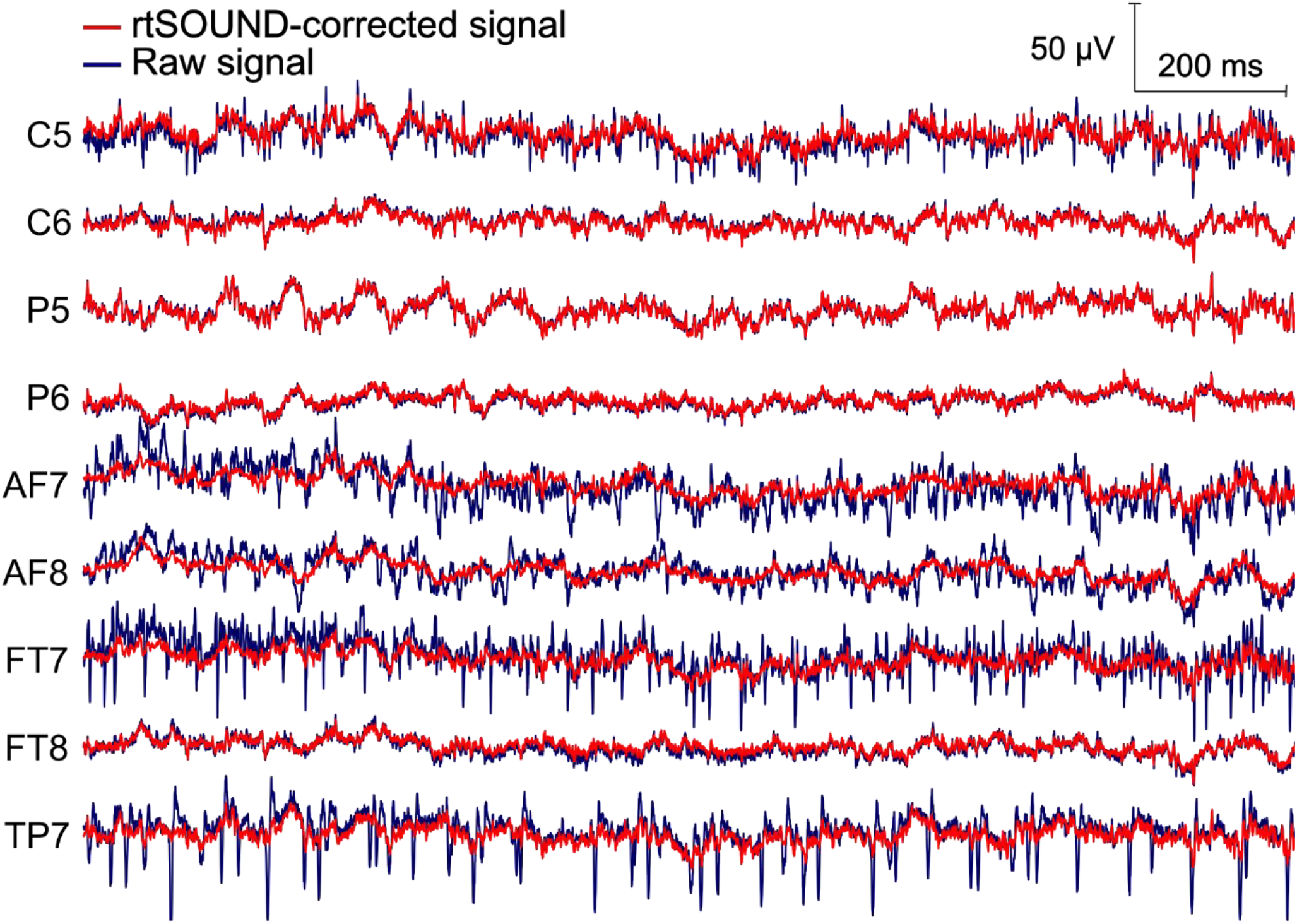
Representative example of real-time baseline-corrected (blue) and rtSOUND-corrected resting-state (red) EEG signals from one of the pre-recorded subjects. The clean signals in the centro-parietal regions (C6, P5, P6) are not altered considerably by rtSOUND. The lateral channels (AF7, AF8, FT7, TP7) have noise likely from muscle activity, which, based on visual inspection, is accurately detected and removed by rtSOUND.

**Fig 5.**
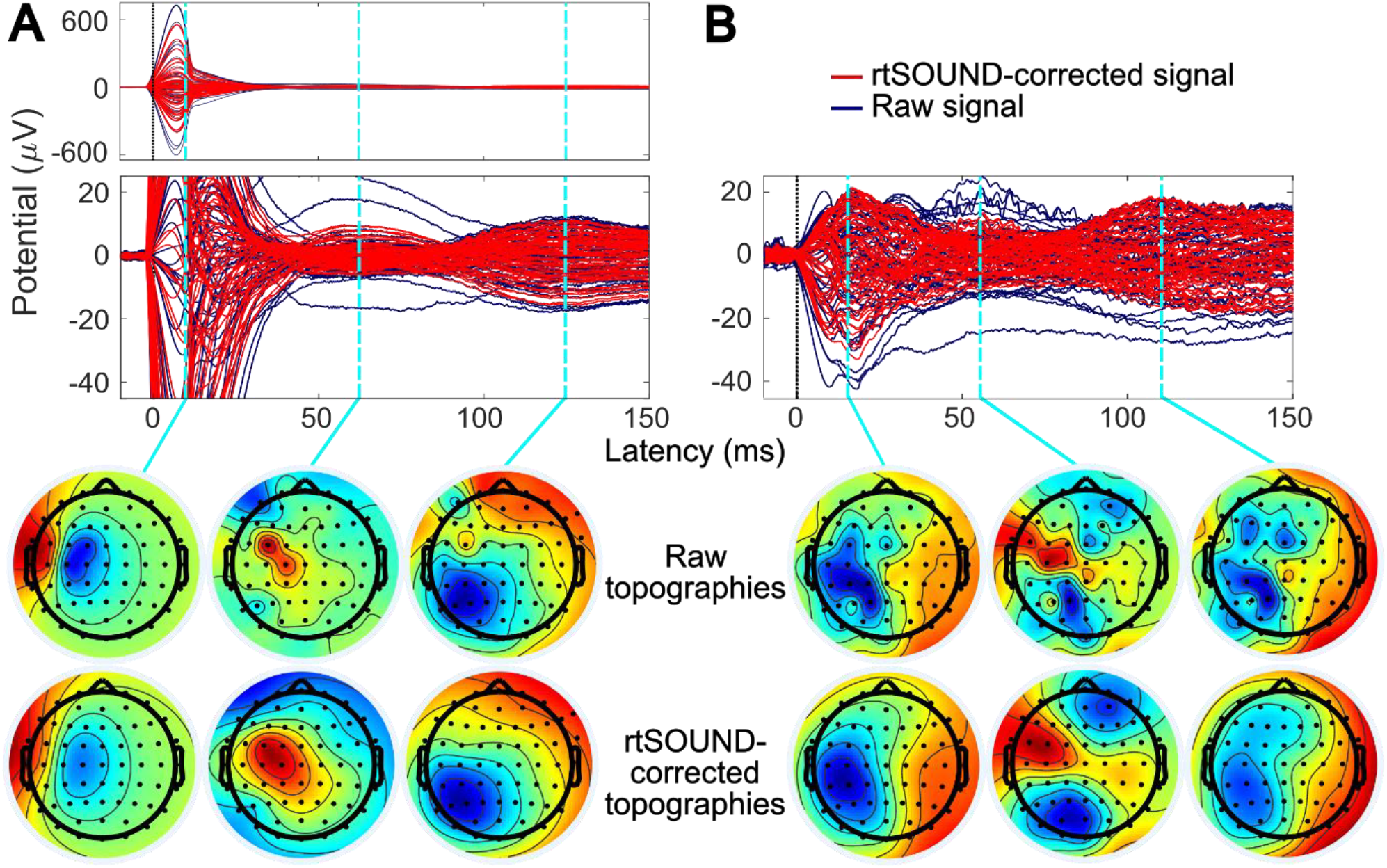
Representative examples of real-time baseline-corrected (blue) and rtSOUND-corrected (red) TMS-evoked potentials from two subjects’ pre-recorded data. Panels A and B represent average TEPs over 70 and 59 epochs, respectively. The topographies of the TEP peaks are also shown; the scale is normalized for each topography to better visualize the changes between baseline-corrected and rtSOUND-corrected signal. In panel A, rtSOUND corrects the noisy channels with large-amplitude fluctuations, revealing a clear response at approximately 60 ms post-pulse. rtSOUND also suppresses, but does not remove, the large TMS-evoked muscle artifact. In panel B, rtSOUND fully corrects the noisy channels. In both panels, rtSOUND smoothens the topographies of the TEP peaks. The TEPs were preprocessed for visualization by removing the baseline (2000 to 10 ms before the pulse), replacing the pulse artifact timespan (–2 to 10 ms) with a cubic interpolation based on data 1 ms before and after the timespan, and rereferencing the data to global average.

#### 3.1.1. SNR changes after rtSOUND

rtSOUND improved SNR of the resting-state data for all 25 subjects. Fig. 6 presents the ICA-based SNR values, which show a wide range of SNR improvements. In the TMS−EEG datasets, SNR was improved for all but one subject, when independent components that were estimated to arise with a probability greater than 0.9 from the brain were labelled as non-artifactual. The overall SNR results were not sensitive to the probability threshold; when it was lowered to 0.8, only one additional subject exhibited a decrease in SNR after rtSOUND. Notably, a qualitative visual inspection of the TEP data for subjects with decreased SNR revealed fewer artifacts post-rtSOUND, despite the metric indicating a reduction in signal quality.

**Fig 6.**
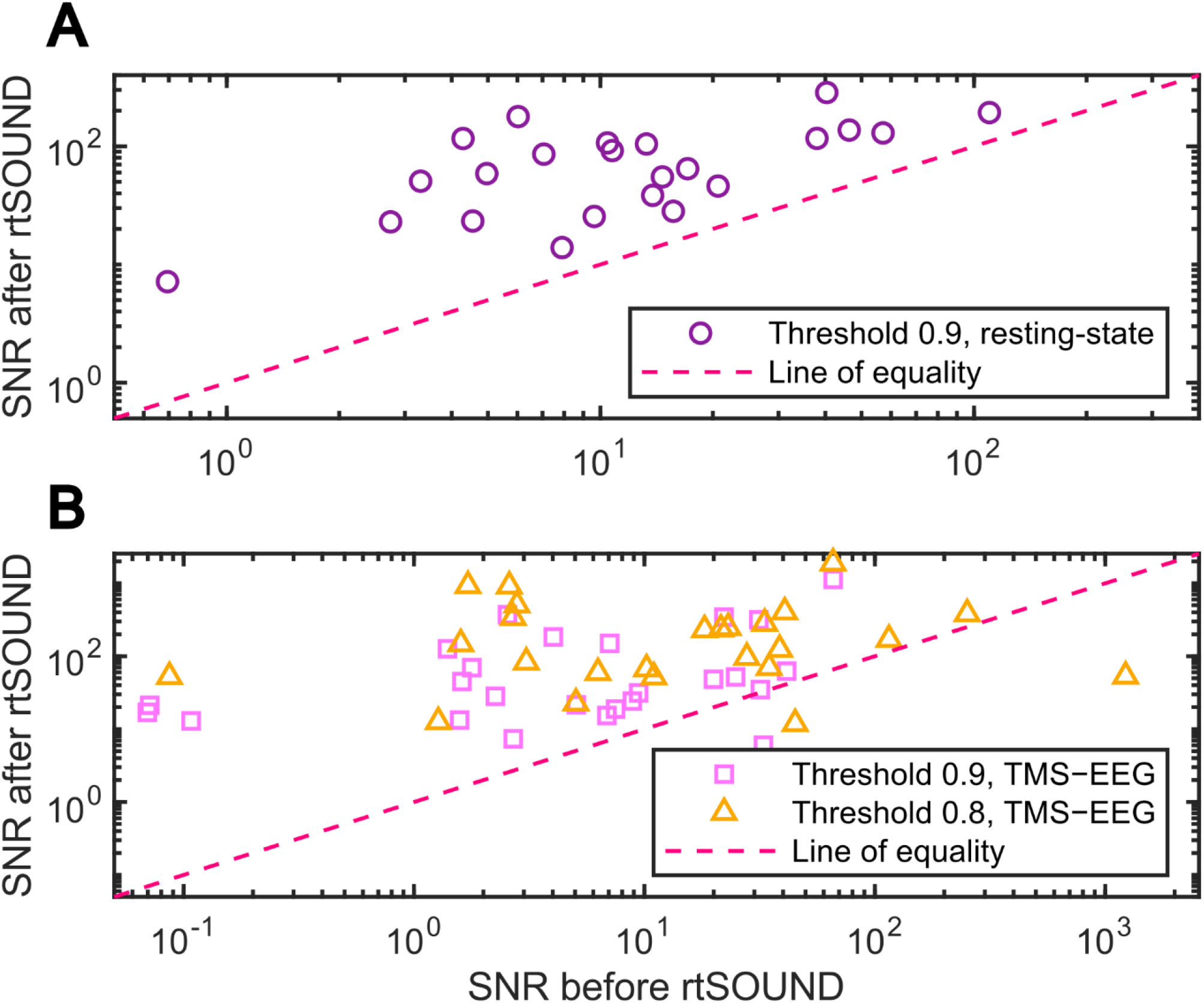
SNR before and after rtSOUND for the resting-state EEG datasets (panel A, *n*=25) with ICA−SNR brain-component labelling threshold 0.9 (purple circles), and TMS−EEG datasets (panel B, *n*=25) with ICA−SNR brain-component labelling thresholds of 0.9 (pink squares) and 0.8 (orange triangles).

#### 3.1.2. Interpolation comparison

The rtSOUND correction produced results very similar to traditional interpolation. The time correlations between the rtSOUND-corrected and traditionally interpolated signals in the three lowest-SNR channels were 0.93 [0.89−0.96] (median and interquartile range over all three channels and trials) for both resting-state and TMS−EEG trials. The relative RMSD were 0.44 [0.32−0.57] for resting-state and 0.49 [0.35−0.72] for TMS−EEG trials. In comparison, the relative RMSD between rtSOUND-corrected and raw signals in the three highest-SNR channels were larger: 0.52 [0.32−0.80] for resting-state and 0.53 [0.31−0.91] for TMS−EEG trials.

#### 3.1.3. Correction of sporadic noise signals: tests with simulation data

In tests with simulated data, rtSOUND began correcting added noise on average 220 ms after the noise onset. However, in some channels and instances, the correction started immediately, suggesting that already the previously updated SOUND filter was tuned to correct the noisy channel. Compared to the noiseless ground-truth data, the relative RMSD decreased by 95% when corrected with rtSOUND.

#### 3.1.4. Relationship between original SNR and change in signal amplitude

LME model analysis provided evidence that the initial SNR has a significant effect on the change in signal amplitude after rtSOUND correction. The LME model ChangeInSignal ∼ 1 + InitialSNR + (1 | Subject) + (1 | Channel) gave a coefficient estimate of −0.35 for initial SNR (*p*-value < 0.001, 95% confidence interval (CI) [−0.42, −0.29]) for the resting-state data. The covariance parameters for the random effects of subjects and channels were estimated to be 0.41 (CI [0.30, 0.55]) and 0.70 (CI [0.58, 0.83]), respectively, implying that the modelled random effects were significant. For the TMS−EEG data, the coefficient estimate for the initial SNR was −0.14 (*p* < 0.001, CI [−0.19, −0.082]). The covariance parameters for the random effects of subjects and channels were estimated to be 0.39 (CI [0.29, 0.52]) and 0.47 (CI [0.39, 0.58]), respectively. The negative coefficient estimates mean that the lower the initial SNR was, the greater was the change in the rtSOUND correction. The results also suggest that both subject-specific and channel-specific random effects need to be considered when modeling the change in signal amplitude.

In addition to the LME results, Spearman correlations between the initial SNR and the change in SNR demonstrated that a lower initial SNR is associated with a larger relative increase in SNR after rtSOUND correction. The correlations were −0.75 (*p* < 0.001) for the resting-state data and −0.72 (*p* < 0.001) for the TMS−EEG datasets.

In the high-quality channels, the time course shapes were preserved, indicated by the correlation analysis. The time correlations between the rtSOUND-corrected and raw signals in the three highest-SNR channels were 0.96 [0.95−0.98] (median and interquartile range over all three channels and trials) for resting-state and 0.93 [0.83−0.97] for TMS−EEG trials. However, as expected [11], rtSOUND exhibited some attenuation in the signal amplitudes even in the best-quality channels. The relative RMSD between rtSOUND-corrected and raw signals in the three highest-SNR channels were 0.52 [0.32−0.80] for resting-state and 0.53 [0.31−0.91] for TMS−EEG trials.

### 3.2. Results of validation with real-time-streamed data

The streamed validation measurements confirmed that rtSOUND performs as intended in real-time settings. The latency measurements showed that both the SOUND filter update and single-sample processing are fast enough for efficient real-time processing and timely adaptation to changing noise parameters. The SOUND filter update duration was typically in the range of 5 to 7 ms, with the observed maximum below 22 ms.

Single-sample processing times were also consistent across measurements, with medians between 61 and 67 μs, and 95^th^ quantiles between 110 and 116 μs, indicating that processing times remained well below the 200-μs threshold required to keep up with the 5-kHz sampling rate, although occasionally individual samples may have taken longer to process. The maximum single-sample processing times ranged from

0.87 to 1.3 ms. SNR of resting-state measurements increased 55–713% after rtSOUND correction.

rtSOUND reduced sporadic noise in less than 150 ms on average. In the four real-time-streamed datasets with added noise, the mean latencies between the onset of noise and rtSOUND’s adaptation ranged from 75 to 133 ms. The adaptation latencies were shorter for subject 2 (mean adaptation time 76 ms) than for subject 1 (mean adaptation time 125 ms).

The power spectra of the signals before and after added noise (Fig. 7) demonstrated that the power distribution of the rtSOUND-corrected signal remained consistent, significantly differing from the noisy raw signal at frequencies above 40 Hz. The relative RMSD between the power spectra of the rtSOUND-corrected signals before and after added noise ranged only from 0.64 [0.54−0.90] to 0.85 [0.75−0.94] (median and interquartile range across noise instances), whereas the corresponding RMSD value between the raw and rtSOUND-corrected noisy signals ranged from 77 [31−610] to 240 [61−600].

**Fig 7.**
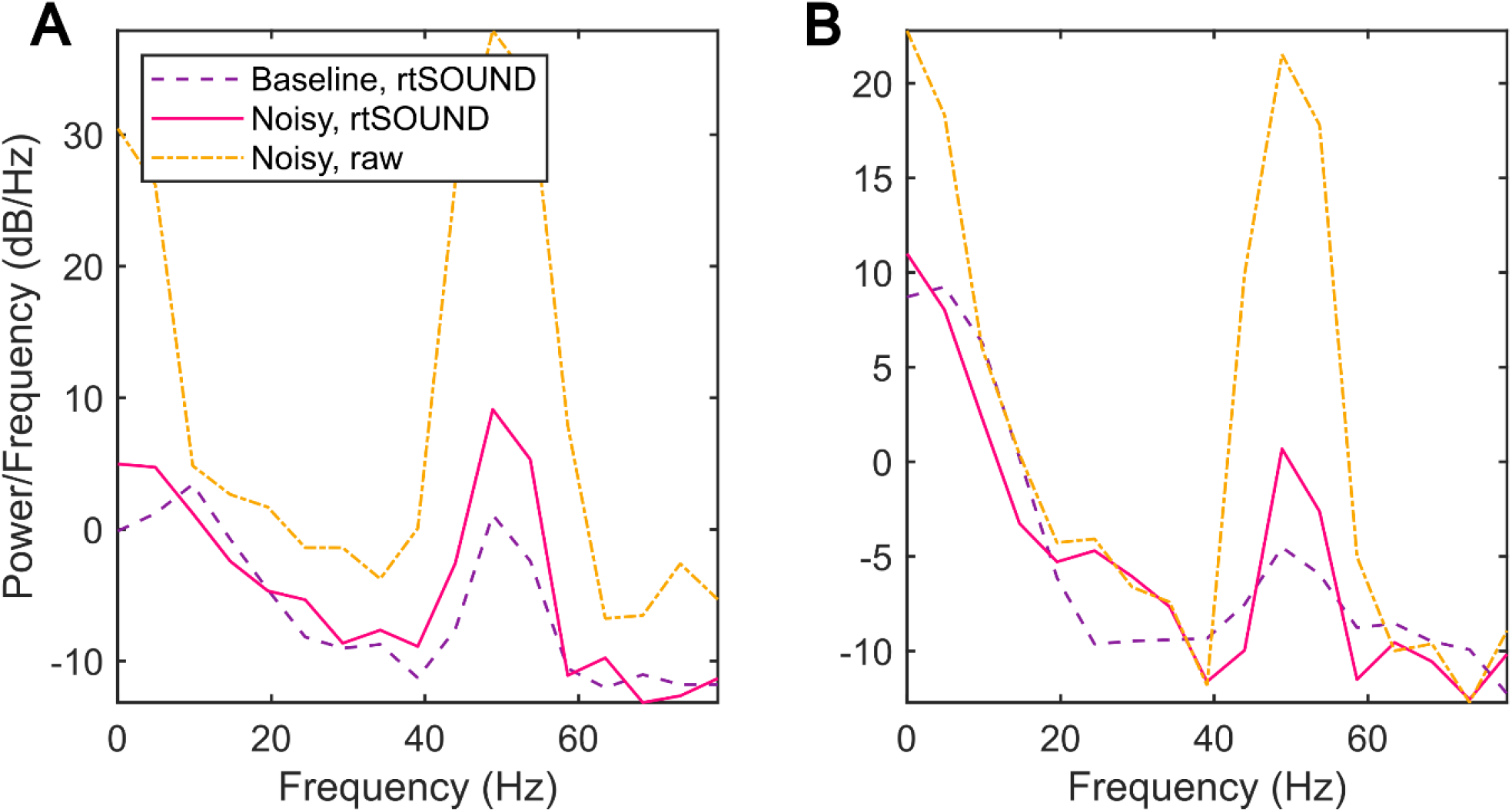
Representative examples of power spectra from one instance of added noise in one channel in the real-time-streamed validation measurements of subject 1 (panel A) and subject 2 (panel B). The spectra represent rtSOUND-corrected data before (purple, dashed) and after (pink, solid) the start of added noise, and the raw (only baseline-corrected) signal after the start of added noise (yellow, dot-dashed). The spectra indicate that rtSOUND effectively suppressed the introduced 50-Hz noise and that the frequency content of the rtSOUND-corrected signals stays similar even when large-amplitude noise is added to the signals.

### 3.3. Added eye-blink artifact-removal results

The results of added eye-blink removal were promising. In 8 out of 10 measurements, SNR was increased by adding blink removal to rtSOUND processing. The median SNR values across the 10 measurements were 22 (min. 11, max. 380) and 61 (min. 14, max. 1000) without and with added blink removal, respectively. Likewise, in 8 out of 10 measurements, the number of eye-artifact components labelled by the automatic IC labelling tool was reduced to zero. Interestingly, in one of the measurements, the number of blink components was not reduced, but SNR was still improved notably. The single-sample processing times with added blink correction were consistent with other validation runs, indicating that blink correction did not slow down the overall processing performance.

### 3.4. Interaction between rtSOUND and TMS-evoked muscle artifacts

Large TMS-evoked muscle artifacts can significantly reduce the achievable post-rtSOUND SNR. The mean post-rtSOUND (pre-rtSOUND) ICA−SNR values were 220

(19) and 36 (8.0) for the TEPs classified into small-(*N*=12) and large-muscle-artifact (*N*=13) groups, respectively. The corresponding median post-rtSOUND (pre-rtSOUND) ICA−SNR values were 100 (14) and 22 (1.8). ICA−SNR between the muscle-artifact groups significantly differed for both post-rtSOUND (*p* = 0.024) and pre-rtSOUND (*p* = 0.018) TEPs. While rtSOUND was able to improve SNR in both large- and small-muscle-artifact groups, the post-rtSOUND SNR values were notably better in the small-than in the large-muscle-artifact group.

### 3.5. Phase effect of rtSOUND

The phase-difference analysis conducted with simulated data showed that rtSOUND can improve phase estimation. The phase differences in channel C3 at the 19-Hz frequency of the simulated resting-state EEG were as follows. Between the ground-truth and noisy data, the circular median of phase differences was 4.3 degrees (range−77 to +70 degrees). Between the ground-truth and rtSOUND-corrected noisy data, the circular median of phase differences was only 1.6 degrees (range −23 to +53 degrees).

## 4. Discussion

We developed a real-time version of the noise-suppressing SOUND algorithm and implemented it into an open-source real-time EEG-processing software. We validated the performance of rtSOUND with pre-recorded resting-state EEG and TMS−EEG datasets from 25 subjects as well as real-time EEG measurements of two subjects. The results indicated that rtSOUND is fast enough to process EEG data streamed at 5 kHz and can improve SNR across the board, as assessed by different metrics. A detailed analysis revealed that rtSOUND primarily modifies channels with poor SNR, suggesting that while it may slightly overcorrect high-quality channels, it still provides a clear overall benefit to EEG signal quality.

### USE CASES OF RTSOUND

rtSOUND has been implemented in the open-source real-time EEG-processing software NeuroSimo [19], facilitating easy deployment into various EEG-based BCI paradigms. Its fully automatic operation enhances usability, requiring the user only to choose the parameters and the lead-field matrix (see section Practical considerations). With the tested parameters, rtSOUND adapts to sporadic noise signals in less than 300 ms, enabling fast SNR improvement during measurements.

rtSOUND can easily be implemented for brain-state-dependent stimulation applications. The current implementation of rtSOUND automatically excludes periods with TMS pulse artifacts or TEPs for updating the noise-suppressing spatial filter, allowing for cleaning of TEPs using the noise distribution estimated from the most recent non-TEP data. However, because the noise distribution during TMS-evoked artifacts likely differs significantly from that of resting-state data, the rtSOUND algorithm should be adapted in the future to separately clean the early tens of milliseconds following each TMS pulse, if the TMS-evoked potentials are of particular interest. In addition, as large TMS-evoked muscle artifacts are correlated between EEG channels and thus violate the assumption of uncorrelated noise, rtSOUND does not fully suppress them (Fig. 5). Large muscle artifacts can also limit SNR achievable by rtSOUND, as shown in our analysis. Thus, while rtSOUND also attenuates muscle artifacts to an extent, obtaining muscle-artifact-free TEPs requires real-time processing methods targeted towards muscle artifacts. In the future, rtSOUND should be tested in combination with other spatial filtering methods, such as signal-space projection–source-informed reconstruction (SSP–SIR) [12, 36].

We compared rtSOUND to a standard EEG cleaning method, interpolation, and found the results to be very similar. However, unlike rtSOUND, interpolation is challenging to perform in real-time due to its time-consuming nature and the need for identifying noisy channels.

The response time of rtSOUND for correcting sporadic noise signals could be improved further by updating the SOUND filter more often than every 100 ms. Based on maximum update durations, updates as frequent as every 30 ms might be feasible, although not tested in this study.

In addition to rtSOUND, several other real-time EEG artifact- and noise-rejection techniques exist. These include Riemannian artifact subspace reconstruction (rASR), an adaptive EOG regression approach, and robust re-referencing. In the present study, we chose not to benchmark rtSOUND directly against these methods, as they are tailored to fundamentally different noise characteristics than those addressed by SOUND. rtSOUND suppresses noise that is uncorrelated across channels, while EOG regression specifically removes ocular artifacts [37, 38]. rASR addresses both ocular and muscle artifacts but requires calibration with artifact-free data [39, 40]. Robust re-referencing mitigates artifacts arising from the reference signal, which are often seen across channels [41-43]. Further studies are needed to examine how these methods can be optimally combined in real-time applications to suppress a wide range of noise and artifact signals.

### PRACTICAL CONSIDERATIONS

The performance of rtSOUND relies on tunable parameters that control its responsiveness, stability, and signal preservation, tailored to the use case and research question. The regularization parameter λ controls noise suppression, with larger values providing greater attenuation but potentially suppressing neuronal signals. A heuristically chosen tuning scalar coefficient λ_0_ is used to calculate λ as follows (see details in the Supplementary material):

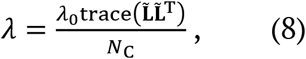

where λ_0_ is the inverse of the estimated SNR. λ_0_ should be tuned to optimize the trade-off between noise removal and overcorrection. In the measurements of this study, we kept λ_0_ fixed at 0.1, which was found optimal in previous studies [11].

The trade-off between rapid adaptation to new noise and stable noise estimates is governed by three parameters: the buffer length, the filter update interval, and the noise-covariance update rate β (see detailed explanation in the Supplementary material). We used a short 100-ms buffer and an equal filter-update interval to assess the responsiveness of rtSOUND to sporadic noise signals. The noise-covariance update rate β was introduced to ensure signal stability and smoothness between consecutive SOUND filters, as large changes in the spatial filter can lead to sudden shifts in the rtSOUND-corrected signal. We used a β value of 0.05. If a swift response to sudden noise is prioritized, the buffer should be short and the SOUND filter update rate high, with β set to minimize shifts in the signal between consecutive filters. However, a short buffer might not capture the characteristics of low-frequency noise in the noise-covariance estimate, risking incomplete removal of long-cycle noise. If the noise is stationary and assumed to change slowly, a longer buffer and filter update interval can be used.

Table S1 in the Supplementary material provides more information about the parameters. An implementation of rtSOUND is available at https://github.com/matildamakkonen/rt-sound-neurosimo-plugin, with the file rtSOUND.py containing the parameters used.

In addition to the parameters, the forward model (lead-field matrix) used with rtSOUND can also affect the cleaning outcome. Compared to source-reconstruction methods, SOUND is more robust against forward model inaccuracies, since the forward model is used only to accurately depict cross-correlations between EEG channels [11, 12]. While SOUND can successfully clean data with a three-layer spherical model or moderate errors in head-tissue conductivity values, inaccurate head geometry can lead to overcorrection, especially in lateral channels [11]. Accordingly, anatomically more accurate forward models improve SOUND performance [11, 12].

We suggest modeling the physical reference channel as part of the forward model. This allows operating the data in its original reference, which is both faster in real-time processing, and decreases the possibility of noise spreading from the reference signal to all channels. In addition, we strongly suggest avoiding reference channel locations that are poorly represented in head models (*e*.*g*., the nose). Instead, we recommend choosing the reference within the International 10–20 or 10–5 channel locations for easier modeling. Theoretical electrode locations should match the actual layout; while digitized locations offer the highest accuracy, they are often impractical as their use requires preparing the digitization and forward model between EEG preparation and the measurement.

### LIMITATIONS

The real-time baseline correction may sometimes induce sudden baseline shifts in the continuous signal. The shifts are minimized with the geometric-sum-based baseline update. Since the baseline correction aims to facilitate real-time analysis, the correction period and frequency can be adjusted to ensure that such shifts do not interfere with the data segment being analyzed (see the Supplementary material for details).

A small previously known disadvantage of SOUND [11] is the slight suppression of amplitudes in the less noisy EEG channels, which is a trade-off for noise removal. The overcorrection level strongly depends on the value of the regularization scalar λ_0_ [11]. However, overcorrection does not alter the topographies of non-noisy signals [11]. SOUND with an optimally chosen regularization scalar has previously been shown to preserve signals well in high-quality channels, especially with high-density sensor layouts [11]. On the other hand, this implies that SOUND is not best suited for EEG applications with a low channel count, such as ICU monitoring.

Overcorrection can be minimized by adaptively changing the value of the regularization scalar λ_0_ during the measurement. This can be useful in long measurements where noise conditions are expected to change, creating the need to adapt the regularization level. An appropriate level of λ_0_ could be determined by estimating the overcorrection in the least noisy channel (based on the noise covariance matrix) after each SOUND filter update. The overcorrection could then be quantified as a relative change between the rtSOUND-corrected and only baseline-corrected EEG buffers. If the relative error exceeds the chosen upper threshold (*e*.*g*., 15%), λ_0_ would be decreased by a learning rate (*e*.*g*., 0.01). Conversely, if the relative error is smaller than the chosen lower threshold (*e*.*g*., 10%), λ_0_ would be increased by the learning rate. Comprehensive tests of this gating mechanism should be performed in future studies.

The phase effects of rtSOUND were studied only offline with a small simulated dataset due to the availability of a true ground-truth signal. While the simulation results were promising, more extensive investigation of the effects of rtSOUND on phase estimation is warranted, particularly in studies where SOUND is used as a preprocessing step for real-time phase-estimation methods. In addition, measuring the performance of rtSOUND with research-question-dependent metrics, such as potential classification accuracy improvement in BCI applications, would provide a task-relevant assessment of signal quality.

The ICA−SNR metric was constructed to account also for those artifacts that overlap with neuronal signals in the frequency domain. The appropriateness of ICA for discerning artifacts from TEP data depends on the trial-to-trial variability of the time course of the artifact component [44, 45]. If the neuronal and artifactual components do not vary sufficiently independently, ICA might include non-artifactual signal contributions in the artifactual components [44, 45], leading to conservative rather than overestimated SNR values in our ICA−SNR analyses. Despite the ICLabel automatic component labeling not being specifically designed for TMS−EEG, it is a part of an established TMS−EEG preprocessing pipeline developed at Stanford University [46].

The added blink-artifact correction can be valuable in applications requiring real-time removal of eye-originating artifacts. However, as this blink removal still requires additional validation, a more established method should be used for blink removal in post-hoc processing.

### FUTURE DIRECTIONS FOR REAL-TIME SPATIAL-FILTER-BASED ARTIFACT REMOVAL

The spatial-filter operation principle of rtSOUND allows for a straightforward integration of other spatial-filter-based EEG-cleaning methods into the pipeline. A variety of other spatial-filter-based EEG-cleaning methods are described in [18]. One such method is the SSP–SIR algorithm [12, 36] for suppressing TMS-induced muscle artifacts. This could enhance real-time TMS–EEG mapping [47-50] or target engagement, as it allows visualizing the early TEP response also when stimulating brain regions vulnerable to muscle artifacts, which are avoided in the current practice [47-50].

In addition to TMS, rtSOUND could be applied to reduce other related stimulation artifacts. For instance, closed-loop amplitude-modulated transcranial alternating current stimulation (AM−tACS) currently suffers from significant artifacts [35, 51] . The tACS artifact masks the neuronal signal but has a local EEG topography [51], which might differ enough from cortically-originated potential patterns for SOUND to suppress this artifact effectively. rtSOUND could be useful for, *e*.*g*., recovering steady-state visually evoked potentials in closed-loop AM−tACS measurements [35].

In addition to EEG, the SOUND algorithm can suppress noise also in MEG, making rtSOUND suitable for MEG-based BCI and other applications. For scenarios requiring very high-resolution real-time measurements, combining MEG and EEG is advantageous, as SOUND can leverage MEG information to enhance EEG noise removal [11].

## 5. Conclusion

We presented the real-time SOUND (rtSOUND) algorithm for robust, fast and adaptive noise removal from EEG. rtSOUND enables utilizing more sophisticated brain−computer-interface and brain-state-dependent stimulation applications, such as novel artificial intelligence-based brain-state detection algorithms, which are often notably sensitive to noise. rtSOUND is available online at https://github.com/matildamakkonen/rt-sound-neurosimo-plugin as a plugin for the open-source NeuroSimo software for brain-state-dependent stimulation and real-time EEG processing.

## CRediT authorship contribution statement

**Matilda Makkonen:** Conceptualization, Data curation, Formal analysis, Funding acquisition, Investigation, Methodology, Software, Validation, Visualization, Writing – original draft, Writing – review and editing. **Olli-Pekka Kahilakoski:** Methodology, Software, Writing – review and editing. **Johanna Metsomaa:** Conceptualization, Methodology, Writing – review and editing. **Pantelis Lioumis:** Funding acquisition, Supervision, Writing – review and editing. **Risto J. Ilmoniemi:** Funding acquisition, Supervision, Writing – review and editing. **Tuomas P. Mutanen:** Conceptualization, Formal analysis, Funding acquisition, Investigation, Methodology, Project administration, Software, Supervision, Validation, Writing – original draft, Writing – review and editing.

## Data availability

De-identified data is available from the corresponding author upon reasonable request, subject to ethical approval and data agreements.

## Code availability

The main code used in this study is publicly available and can be accessed at https://github.com/matildamakkonen/rt-sound-neurosimo-plugin.

## Declaration of competing interest

For this work, TPM received funding for a collaborative research project with Bittium Biosignals Oy (Kuopio, Finland). RJI is a founder of Cortisys Inc. PL is a consultant to Nexstim Plc. for TMS–EEG applications and speech cortical mapping.

### Declaration of generative AI and AI-assisted technologies in the writing process

During the preparation of this work the authors used ChatGPT-3.5 and ChatGPT-4o in order to edit and proofread the language of the raw text. After using this tool, the authors reviewed and edited the content as needed and take full responsibility for the content of the published article.

## Acknowledgements

We thank the researchers involved in collecting the 25-subject dataset: Oskari Ahola, Mikael Laine, Noora Kainulainen, Joonas Laurinoja, Ana Soto de la Cruz, Shokoofeh Parvin, Maria Nazarova, Ida Granö, Mikko Nyrhinen, Ilkka Rissanen, Ida Ilmoniemi, and Elena Ukharova. We also thank Aura Ruokoranta for the help with the correlation analysis. This work was supported by the Emil Aaltonen Foundation (MM; grant number 240110 K1), the Aalto University US Pilot Program (MM), the European Research Council (ERC Synergy) under the European Union’s Horizon 2020 research and innovation programme (ConnectToBrain; grant agreement No 810377), and the Finnish Foundation for Technology Promotion (TPM).

## References

[1] Tatum WO, Rubboli G, Kaplan PW, Mirsatari S, Radhakrishnan K, Gloss D, et al. Clinical utility of EEG in diagnosing and monitoring epilepsy in adults. Clinical Neurophysiology 2018;129(5):1056–82.

[2] Rundo JV, Downey III R. Polysomnography. Handbook of clinical neurology 2019;160:381–92.

[3] Värbu K, Muhammad N, Muhammad Y. Past, present, and future of EEG-based BCI applications. Sensors 2022;22(9):3331.

[4] Cao Z. A review of artificial intelligence for EEG-based brain− computer interfaces and applications. Brain Science Advances 2020;6(3):162–70.

[5] Zubarev I, Zetter R, Halme H-L, Parkkonen L. Adaptive neural network classifier for decoding MEG signals. NeuroImage 2019;197:425–34.

[6] Minguillon J, Lopez-Gordo MA, Pelayo F. Trends in EEG-BCI for daily-life: Requirements for artifact removal. Biomedical Signal Processing and Control 2017;31:407–18.

[7] Ferree TC, Luu P, Russell GS, Tucker DM. Scalp electrode impedance, infection risk, and EEG data quality. Clinical neurophysiology 2001;112(3):536–44.

[8] Cohen L. Time-Frequency Analysis. Upper Saddle River, New Jersey: Prentice Hall PTR; 1995.

[9] Chen LL, Madhavan R, Rapoport BI, Anderson WS. Real-time brain oscillation detection and phase-locked stimulation using autoregressive spectral estimation and timeseries forward prediction. IEEE transactions on Biomedical Engineering 2011;60(3):753–62.

[10] Zrenner C, Desideri D, Belardinelli P, Ziemann U. Real-time EEG-defined excitability states determine efficacy of TMS-induced plasticity in human motor cortex. Brain stimulation 2018;11(2):374–89.

[11] Mutanen TP, Metsomaa J, Liljander S, Ilmoniemi RJ. Automatic and robust noise suppression in EEG and MEG: The SOUND algorithm. Neuroimage 2018;166:135–51.

[12] Mutanen TP, Metsomaa J, Makkonen M, Varone G, Marzetti L, Ilmoniemi RJ. Source-based artifact-rejection techniques for TMS–EEG. Journal of Neuroscience Methods 2022;382:109693.

[13] Berger H. ^Ü^ber das elektroenkephalogramm des menschen. Archiv für psychiatrie und nervenkrankheiten 1929;87(1):527–70.

[14] Niedermeyer E, da Silva FL. Electroencephalography: basic principles, clinical applications, and related fields. Lippincott Williams & Wilkins; 2005.

[15] Khanna A, Pascual-Leone A, Michel CM, Farzan F. Microstates in resting-state EEG: current status and future directions. Neuroscience & Biobehavioral Reviews 2015;49:105–13.

[16] Hernandez-Pavon JC, Veniero D, Bergmann TO, Belardinelli P, Bortoletto M, Casarotto S, et al. TMS combined with EEG: Recommendations and open issues for data collection and analysis. Brain Stimulation 2023;16(2):567–93.

[17] Weinstein D, Zhukov L, Johnson C. Lead-field bases for electroencephalography source imaging. Annals of biomedical engineering 2000;28:1059–65.

[18] Metsomaa J, Song Y, Mutanen TP, Gordon PC, Ziemann U, Zrenner C, et al. Adapted Beamforming: A Robust and Flexible Approach for Removing Various Types of Artifacts from TMS–EEG Data. Brain Topography 2024;37(5):659–83.

[19] Kahilakoski O-P, Alkio K, Siljamo O, Valén K, Laurinoja J, Haxel L, et al. NeuroSimo: an open-source software for closed-loop EEG-or EMG-guided TMS. bioRxiv 2025:2025.04. 05.647342.

[20] Macenski S, Foote T, Gerkey B, Lalancette C, Woodall W. Robot operating system 2: Design, architecture, and uses in the wild. Science robotics 2022;7(66):eabm6074.

[21] Koponen LM, Nieminen JO, Ilmoniemi RJ. Multi-locus transcranial magnetic stimulation—theory and implementation. Brain Stimulation 2018;11(4):849–55.

[22] Nieminen JO, Sinisalo H, Souza VH, Malmi M, Yuryev M, Tervo AE, et al. Multi-locus transcranial magnetic stimulation system for electronically targeted brain stimulation. Brain Stimulation 2022;15(1):116–24.

[23] Huang Y, Parra LC, Haufe S. The New York Head—A precise standardized volume conductor model for EEG source localization and tES targeting. NeuroImage 2016;140:150–62.

[24] Ziemann U, Romani G-L, Ilmoniemi RJ. ‘ConnectToBrain’: Synergy-Projekt zur Closed-loop-Stimulationstherapie von Netzwerkerkrankungen des Gehirns. Der Nervenarzt 2019;90(8):p804.

[25] Russo S, Sarasso S, Puglisi GE, Dal Palù D, Pigorini A, Casarotto S, et al. TAAC-TMS Adaptable Auditory Control: A universal tool to mask TMS clicks. Journal of neuroscience methods 2022;370:109491.

[26] Rocchi L, Di Santo A, Brown K, Ibánez J, Casula E, Rawji V, et al. Disentangling EEG responses to TMS due to cortical and peripheral activations. Brain stimulation 2021;14(1):4–18.

[27] Pion-Tonachini L, Kreutz-Delgado K, Makeig S. ICLabel: An automated electroencephalographic independent component classifier, dataset, and website. NeuroImage 2019;198:181–97.

[28] Hyvärinen A, Oja E. Independent component analysis: algorithms and applications. Neural networks 2000;13(4-5):411–30.

[29] Perrin F, Pernier J, Bertrand O, Echallier JF. Spherical splines for scalp potential and current density mapping. Electroencephalography and clinical neurophysiology 1989;72(2):184–7.

[30] Delorme A, Makeig S. EEGLAB: an open source toolbox for analysis of single-trial EEG dynamics including independent component analysis. Journal of neuroscience methods 2004;134(1):9–21.

[31] Barr DJ, Levy R, Scheepers C, Tily HJ. Random effects structure for confirmatory hypothesis testing: Keep it maximal. Journal of memory and language 2013;68(3):255–78.

[32] Makkonen M, Mutanen T, Metsomaa J, Zrenner C, Souza V, Ilmoniemi R. Real-time artifact detection and removal for closed-loop EEG-TMS. International Journal of Bioelectromagnetism 2021;23(2):1–4.

[33] Lehmann D, Skrandies W. Reference-free identification of components of checkerboard-evoked multichannel potential fields. Electroencephalography and clinical neurophysiology 1980;48(6):609–21.

[34] Wilcoxon F. Individual comparisons by ranking methods. Biometrics bulletin 1945;1(6):80–3.

[35] Haslacher D, Cavallo A, Reber P, Kattein A, Thiele M, Nasr K, et al. Working memory enhancement using real-time phase-tuned transcranial alternating current stimulation. Brain Stimulation 2024;17(4):850–9.

[36] Mutanen TP, Kukkonen M, Nieminen JO, Stenroos M, Sarvas J, Ilmoniemi RJ. Recovering TMS-evoked EEG responses masked by muscle artifacts. Neuroimage 2016;139:157–66.

[37] He P, Wilson G, Russell C. Removal of ocular artifacts from electro-encephalogram by adaptive filtering. Medical and biological engineering and computing 2004;42(3):407–12.

[38] He P, Wilson G, Russell C, Gerschutz M. Removal of ocular artifacts from the EEG: a comparison between time-domain regression method and adaptive filtering method using simulated data. Medical & biological engineering & computing 2007;45(5):495–503.

[39] Blum S, Jacobsen NS, Bleichner MG, Debener S. A Riemannian modification of artifact subspace reconstruction for EEG artifact handling. Frontiers in human neuroscience 2019;13:141.

[40] Arpaia P, De Bendetto E, Esposito A, Natalizio A, Parvis M, Pesola M. Comparing artifact removal techniques for daily-life electroencephalography with few channels. 2022 IEEE International Symposium on Medical Measurements and Applications (MeMeA). IEEE; 2022:1–6.

[41] Lepage KQ, Kramer MA, Chu CJ. A statistically robust EEG re-referencing procedure to mitigate reference effect. Journal of neuroscience methods 2014;235:101–16.

[42] Bigdely-Shamlo N, Mullen T, Kothe C, Su K-M, Robbins KA. The PREP pipeline: standardized preprocessing for large-scale EEG analysis. Frontiers in neuroinformatics 2015;9:16.

[43] de Cheveigné A, Arzounian D. Robust detrending, rereferencing, outlier detection, and inpainting for multichannel data. NeuroImage 2018;172:903–12.

[44] Atti I, Belardinelli P, Ilmoniemi RJ, Metsomaa J. Measuring the accuracy of ICA-based artifact removal from TMS-evoked potentials. Brain stimulation 2024;17(1):10–8.

[45] Mutanen TP, Ilmoniemi I, Atti I, Metsomaa J, Ilmoniemi RJ. A simulation study: comparing independent component analysis and signal-space projection–source-informed reconstruction for rejecting muscle artifacts evoked by transcranial magnetic stimulation. Frontiers in Human Neuroscience 2024;18:1324958.

[46] Cline CC, Lucas MV, Sun Y, Menezes M, Etkin A. Advanced artifact removal for automated TMS-EEG data processing. 2021 10th International IEEE/EMBS Conference on Neural Engineering (NER). IEEE; 2021:1039–42.

[47] Casarotto S, Fecchio M, Rosanova M, Varone G, D’Ambrosio S, Sarasso S, et al. The rt-TEP tool: real-time visualization of TMS-Evoked Potentials to maximize cortical activation and minimize artifacts. Journal of Neuroscience Methods 2022;370:109486.

[48] Parmigiani S, Cline CC, Sarkar M, Forman L, Truong J, Ross JM, et al. Real-time optimization to enhance noninvasive cortical excitability assessment in the human dorsolateral prefrontal cortex. Clinical Neurophysiology 2025;174:225–34.

[49] Lioumis P, Rosanova M. The role of neuronavigation in TMS–EEG studies: Current applications and future perspectives. Journal of Neuroscience Methods 2022;380:109677.

[50] Ukharova E, Sathyan S, GranöI, O’ Meeghan I, Ahola O, Kainulainen N, et al. Neuroimaging-Guided TMS–EEG for Real-Time Cortical Network Mapping. Journal of Visualized Experiments (JoVE) 2025(220):e67339.

[51] Haslacher D, Nasr K, Robinson SE, Braun C, Soekadar SR. Stimulation artifact source separation (SASS) for assessing electric brain oscillations during transcranial alternating current stimulation (tACS). Neuroimage 2021;228:117571.

