## Supplementary Material for "Real-time noise cancellation for EEG monitoring and brain–computer interfaces"

### 1. Implementing real-time SOUND algorithm

Our real-time noise-cancellation method is based on the SOurce-estimate-Utilizing Noise-Discarding (SOUND) algorithm [1, 2], which was first developed for offline cleaning of electroencephalography (EEG) [1]. In this supplementary material, we cover the foundations of SOUND and its real-time adaptation in detail.

The forward problem of EEG is to solve the voltage patterns at the scalp resulting from theoretical unit-current dipole sources on the cortex. Each current dipole models local synchronous post-synaptic currents at the defined cortical location. The solution of the forward problem is the so-called lead-field matrix $\mathbf{L}$ [3], whose columns correspond to the voltage patterns (or EEG topographies), generated by individual dipolar current sources.

Once the forward problem has been solved, we can write the measured EEG data in terms of unknown brain-activity distribution. Streaming EEG can be thought of as a measure of topographies $\mathbf{y}$ as a function of time:

$$\mathbf{y}(t) = \begin{bmatrix} y_1(t) \\ \vdots \\ y_{N_C}(t) \end{bmatrix}, \qquad \text{(S1)}$$

where $\mathbf{y}(t)$ is an $N_C \times 1$ signal vector, whose element $y_i(t)$ corresponds to the voltage measured by channel $i$ at time $t$, and $N_C$ is the number of EEG channels. The measured signal $\mathbf{y}$ is the sum of the voltage patterns resulting from active cortical current sources and noise:

$$\mathbf{y}(t) = \mathbf{L}\mathbf{x}(t) + \mathbf{n}(t) = \mathbf{L}\mathbf{x}(t) + \mathbf{M}_{\mathrm{N}}\mathbf{s}_{\mathrm{N}}(t)\ , \quad \text{(S2)}$$

where $\mathbf{L}$ is the $N_{\mathrm{C}} \times N_{\mathrm{S}}$ lead-field matrix ($N_{\mathrm{S}}$ being the number of modelled cortical current sources), $\mathbf{x}(t)$ is an unknown $N_{\mathrm{S}} \times 1$ vector describing the active cortical current distribution at time $t$, and $\mathbf{n}(t)$ is the noise at time $t$. $\mathbf{n}(t)$ can be written in terms of $\mathbf{M}_{\mathrm{N}}$ and $\mathbf{s}_{\mathrm{N}}$ , which are the $N_{\mathrm{C}} \times N_{\mathrm{N}}$ noise mixing matrix and $N_{\mathrm{N}} \times 1$ noise signal vector, respectively. In general, the number of noise components, $N_{\mathrm{N}}$ , is unknown. $\mathbf{M}_{\mathrm{N}}$ holds the topographies of individual noise components in its columns, whereas the rows of $\mathbf{s}_{\mathrm{N}}$ correspond to the time courses of different noise components.

Our objective is to find an $N_{\mathrm{C}} \times N_{\mathrm{C}}$ spatial filter matrix $\mathbf{W}_{\mathrm{SOUND}}$ that suppresses the noise topographies but not the neuronal ones, approximately satisfying $\mathbf{W}_{\mathrm{SOUND}}\mathbf{L} \approx \mathbf{L}$ and $\mathbf{W}_{\mathrm{SOUND}}\mathbf{M}_{\mathrm{N}} \approx 0$. Then

$$\hat{\mathbf{y}}_{\mathrm{clean}}(t) = \mathbf{W}_{\mathrm{SOUND}}\mathbf{y}(t) = \mathbf{W}_{\mathrm{SOUND}}\mathbf{L}\mathbf{x}(t) + \mathbf{W}_{\mathrm{SOUND}}\,\mathbf{M}_{\mathrm{N}}\mathbf{s}_{\mathrm{N}}(t) \approx \mathbf{L}\mathbf{x}(t)\quad . \quad \text{(S3)}$$

To solve for $\mathbf{W}_{\mathrm{SOUND}}$, we first examine the inverse problem of EEG, which refers to solving the unknown cortical currents based on the measurement $\mathbf{y}(t)$ and the lead-field matrix $\mathbf{L}$. Because the problem is severely underdetermined, we must apply generalized inverse techniques to estimate $\mathbf{x}$. A common approach is to compute the minimum-norm estimate (MNE) [4]. For data whitened with respect to the noise,

$$\mathbf{C}_{\mathrm{N}}^{-1/2}\mathbf{y} = \mathbf{C}_{\mathrm{N}}^{-1/2}\mathbf{L}\mathbf{x} + \mathbf{C}_{\mathrm{N}}^{-1/2}\mathbf{n}\ , \qquad \text{(S4)}$$

MNE can be written [5]:

$$\hat{\mathbf{x}} = \tilde{\mathbf{L}}^{\mathrm{T}}(\tilde{\mathbf{L}}\tilde{\mathbf{L}}^{\mathrm{T}} + \lambda\mathbf{I})^{-1}\mathbf{C}_{\mathrm{N}}^{-1/2}\mathbf{y} = \mathbf{K}_{\mathrm{MNE}}\mathbf{y}\ , \qquad \text{(S5)}$$

where $\mathbf{C}_{\mathrm{N}}$ is the $N_{\mathrm{C}} \times N_{\mathrm{C}}$ noise covariance matrix, $\lambda$ is the regularization parameter, $\tilde{\mathbf{L}} = \mathbf{C}_{\mathrm{N}}^{-1/2}\mathbf{L}$ is the whitened version of the lead-field matrix, and $\mathbf{K}_{\mathrm{MNE}} = \tilde{\mathbf{L}}^{\mathrm{T}}(\tilde{\mathbf{L}}\tilde{\mathbf{L}}^{\mathrm{T}} + \lambda\mathbf{I})^{-1}\mathbf{C}_{\mathrm{N}}^{-1/2}$ is the combined MNE–whitening operator. The whitening with respect to the noise can be interpreted as a penalty term in the EEG signal space, based on the structure of the noise; those signal-space directions in which the noise is more pronounced will be suppressed more.

The regularization parameter $\lambda$ defines how much we suppress noise in the brain-activity estimate. After whitening, $\lambda$ can be written [5]:

$$\lambda = \frac{\lambda_0 \mathrm{trace}(\tilde{\mathbf{L}}\tilde{\mathbf{L}}^{\mathrm{T}})}{N_{\mathrm{C}}}, \qquad \text{(S6)}$$

where $\lambda_0$ is the inverse of the estimated SNR.

If the regularization in the MNE operator has successfully attenuated the noise in the source estimates, we can compute noise-suppressed EEG signals from the noise-suppressed brain estimates by multiplying the source estimates with $\mathbf{L}$ from the left:

$$\hat{\mathbf{y}} = \mathbf{L}\hat{\mathbf{x}}\ . \qquad \text{(S7)}$$

By combining Eqs. S5 and S7, we can write the cleaned EEG signals in terms of the original signals, and define the spatial SOUND filter accordingly:

$$\hat{\mathbf{y}} = \mathbf{L}\mathbf{K}_{\mathrm{MNE}}\mathbf{y}$$
$$\Rightarrow \mathbf{W}_{\mathrm{SOUND}}(\mathbf{L}, \mathbf{C}_{\mathrm{N}}, \lambda) = \mathbf{L}\mathbf{K}_{\mathrm{MNE}} \qquad \text{(S8)}$$

From Eq. S8 we can see that to clean the data, we need an estimate for the noise-covariance matrix $\mathbf{C}_{\mathrm{N}}$. If we assume $\mathbf{C}_{\mathrm{N}}$ to be diagonal, its estimate can be written:

$$\hat{\mathbf{C}}_{\mathrm{N}} = \begin{bmatrix} \hat{\sigma}_1^2 & \cdots & 0 \\ \vdots & \ddots & \vdots \\ 0 & \cdots & \hat{\sigma}_{N_{\mathrm{C}}}^2 \end{bmatrix}, \qquad \text{(S9)}$$

where $\hat{\sigma}_i$ is the estimated noise level in channel $i$.

We next describe an efficient method for estimating the channel-specific noise levels, and thus $\hat{\mathbf{C}}_{\mathrm{N}}$. Metsomaa *et al.* [6] recently showed that noise signal can be estimated efficiently with beamforming:

$$S_{\mathrm{N}} = \mathbf{w}_{\mathrm{N}}^{\mathrm{T}}\mathbf{Y}\ , \qquad \text{(S10)}$$

where $S_{\mathrm{N}}$ is the time course of a noise signal N, $\mathbf{w}_{\mathrm{N}}$ is an $N_{\mathrm{C}} \times 1$ spatial filter vector for estimating the noise component, and $\mathbf{Y}$ is the $N_{\mathrm{C}} \times N_{\mathrm{T}}$ data matrix ($N_{\mathrm{T}}$ is the number of time points). Thus, the variance of a noise source, for instance in a noisy channel $i$, can be computed from the data covariance matrix $\mathbf{C}_{\mathrm{Y}}$:

$$\hat{\sigma}_i = \mathbf{w}_i^{\mathrm{T}}\mathbf{C}_{\mathrm{Y}}\mathbf{w}_i\ . \text{ (S11)}$$

If we assume that the noise in each channel is uncorrelated with that of the other channels, we can derive the following formula for the channel-specific spatial filters (see the detailed derivation in [6]):

$$\mathbf{w}_i = \frac{\boldsymbol{\Gamma}_i}{\Gamma_{i,i}}, \text{(S12)}$$

where $\boldsymbol{\Gamma} = (\mathbf{L}\mathbf{L}^{\mathrm{T}} + \lambda \mathbf{C}_{\mathrm{N}})^{-1}$ and $\boldsymbol{\Gamma}_i$ denotes the $i$'th column and $\Gamma_{i,i}$ the element on the $i$'th row and column of $\boldsymbol{\Gamma}$. By combining Eqs. S9, S11, and S12, the full noise covariance matrix estimate $\hat{\mathbf{C}}_{\mathrm{N}}$ can be retrieved efficiently from the data covariance matrix $\mathbf{C}_{\mathrm{Y}}$:

$$\hat{\mathbf{C}}_{\mathrm{N}} = \begin{bmatrix} \left(\frac{\boldsymbol{\Gamma}_i}{\Gamma_{i,i}}\right)^T \mathbf{C}_{\mathrm{Y}} \frac{\boldsymbol{\Gamma}_i}{\Gamma_{i,i}} & \cdots & 0 \\ \vdots & \ddots & \vdots \\ 0 & \cdots & \left(\frac{\boldsymbol{\Gamma}_{N_{\mathrm{C}}}}{\Gamma_{N_{\mathrm{C}},N_{\mathrm{C}}}}\right)^T \mathbf{C}_{\mathrm{Y}} \frac{\boldsymbol{\Gamma}_{N_{\mathrm{C}}}}{\Gamma_{N_{\mathrm{C}},N_{\mathrm{C}}}} \end{bmatrix}. \quad \text{(S13)}$$

Because $\boldsymbol{\Gamma}$ depends on the estimated noise-covariance matrix, which in turn can be estimated by applying channel-wise operators $\frac{\boldsymbol{\Gamma}_i}{\Gamma_{i,i}}$ to the data covariance matrix, we update $\hat{\mathbf{C}}_{\mathrm{N}}$ iteratively until convergence. We can measure the convergence level, *e.g.*, as the maximum relative difference in channel-specific noise variances (diagonal elements of $\hat{\mathbf{C}}_{\mathrm{N}}$) between consecutive iterations.

To use the standard SOUND algorithm for *post-hoc* EEG analysis, one can update the noise covariance matrix with Eq. S13 and clean the data with Eq. S8. Next, we discuss how the SOUND algorithm was adapted to process streaming EEG data in real-time (hereinafter referred to as rtSOUND). The adaptation requires organizing the updating of the noise-covariance estimate, and the application of the resulting SOUND filter to data samples. We pursue this with a simple parallelization scheme.

At time $t_0$, we clean the data sample with the latest SOUND filter estimate, $\widehat{\mathbf{W}}_{\mathrm{SOUND}}(\mathbf{L}, \hat{\mathbf{C}}_{\mathrm{N}}, \lambda)$:

$$\hat{\mathbf{y}}(t_0) = \widehat{\mathbf{W}}_{\mathrm{SOUND}} \mathbf{y}(t_0), \text{(S14)}$$

If no SOUND filter update is ongoing, we launch a new estimation of the noise-covariance matrix. Using Eq. S13 requires the data covariance matrix, $\mathbf{C}_{\mathrm{Y}}$, which we estimate as the sample covariance matrix by fetching the $N$ past samples from the data buffer:

$$\hat{\mathbf{C}}_{\mathrm{Y}} \approx \frac{1}{N-1} \mathbf{Y}(t_{-N+1}, t_{-N+2}, \cdots, t_{-1}, t_0) \left(\mathbf{Y}(t_{-N+1}, t_{-N+2}, \cdots, t_{-1}, t_0)\right)^{\mathrm{T}}. \text{(S15)}$$

Based on Eqs. S8, S13, and S15, we can compute the updated $\widehat{\mathbf{W}}'_{\text{SOUND}}$ and start cleaning the data with the latest filter:

$$\hat{\mathbf{y}}(t_0 + \Delta t) = \widehat{\mathbf{W}}'_{\text{SOUND}}\mathbf{y}(t_0 + \Delta t) \text{ , (S16)}$$

where $\Delta t$ is the update time. Each sample is cleaned separately as soon as it is streamed by the EEG device.

Large changes in consecutive SOUND filters might lead to sudden shifts in the rtSOUND-corrected signal. To mitigate this, we implemented a smooth filter update using a geometric-sum-based update rule for the noise covariance matrix $\hat{\mathbf{C}}_{\text{N}}$. We define an update rate $\beta \in [0,1]$, and after calculating $\hat{\mathbf{C}}_{\text{N}}^{\text{new}}$ based on the most recent data buffer, we utilize the noise covariance matrix $\hat{\mathbf{C}}_{\text{N}}^{\text{prev}}$ from the previous SOUND update to compute $\hat{\mathbf{C}}_{\text{N}}^{\text{smooth}}$ for updating the SOUND filter:

$$\hat{\mathbf{C}}_{\text{N}}^{\text{smooth}} = \beta\hat{\mathbf{C}}_{\text{N}}^{\text{new}} + (1-\beta)\hat{\mathbf{C}}_{\text{N}}^{\text{prev}} \text{ . (S17)}$$

For SOUND to reliably estimate the true channel-specific noise levels, and hence, to filter the data accurately, it is critical that the data are baseline-corrected. We can apply a simple baseline-correction algorithm to the streaming data. Let

$$\mathbf{y}(t_0) = \mathbf{y}^{\text{r}}(t_0) - \mathbf{b}(t_0) \text{ , (S18)}$$

where $\mathbf{y}^{\text{r}}(t_0)$ is an $N_{\text{C}} \times 1$ vector holding the raw voltage values coming directly from an EEG amplifier operating in direct current (DC) mode, at time $t_0$, and $\mathbf{b}(t_0)$ is an $N_{\text{C}} \times 1$ time-adaptive baseline correction vector defined as

$$\mathbf{b}(t_0) = \alpha\mathbf{y}^{\text{r}}(t_0) + (1-\alpha)\mathbf{b}(t_{-1}) \text{ , (S19)}$$

where $\alpha \in [0,1]$ is the update rate of the baseline correction. Note that for an EEG amplifier operating in alternating current (AC) mode, the baseline correction is not necessary. The whole workflow of running rtSOUND is schematically visualized in Fig. S1.

As described in the mathematical formulation of the algorithm above, the performance of rtSOUND depends on several tunable parameters that govern its responsiveness, stability, and signal preservation. Table S1 summarizes the specific parameter values used in this study, along with their intended purpose and the rationale behind each choice. Overall, these values were selected to ensure efficient noise suppression while maintaining the integrity of the underlying EEG signal in real-time applications. In addition to Table S1, the effects of different update rates on baseline-correction behavior are visualized in Fig. S2.

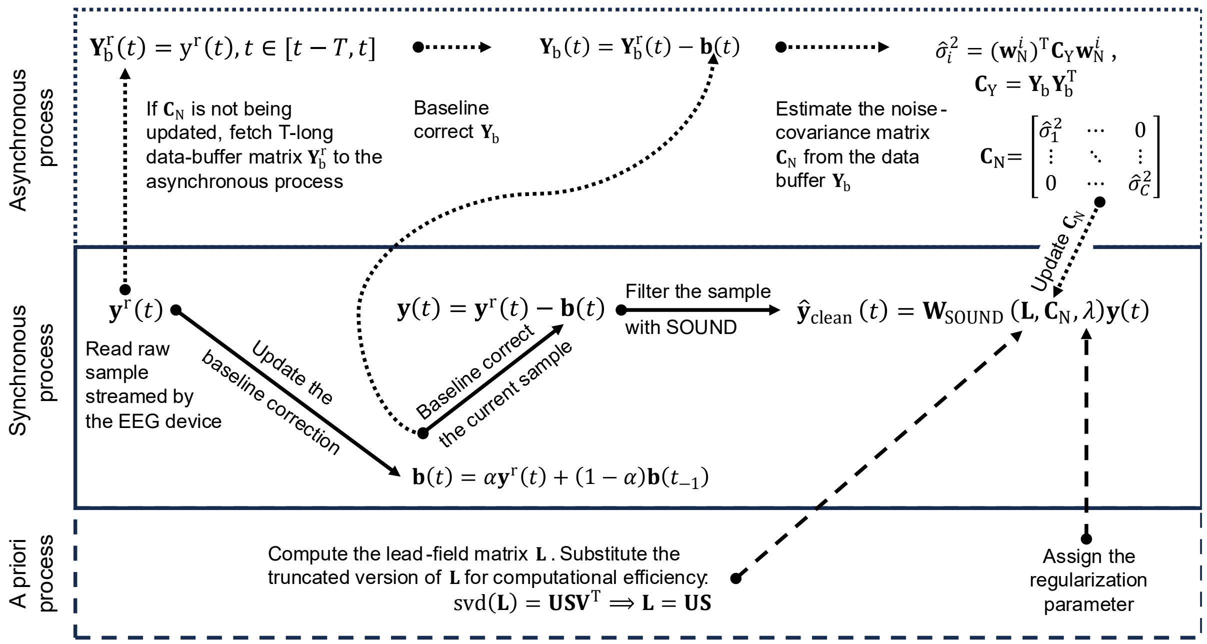


**Fig S1.** Summary of the real-time noise-cancellation algorithm rtSOUND. Before the measurement, the user provides a suitable lead-field matrix and a value of $\lambda$, dictating the regularization level (Eq. S6). Note that we use the truncated version of the lead-field matrix for computational efficiency of the pipeline [2]. When the measurement begins, each sample of raw data is processed by rtSOUND in a real-time (synchronous) process. For each sample, the baseline correction is first updated according to Eq. S19, and then the baseline is deducted from the sample. The clean sample is obtained by multiplying the sample with the latest SOUND filter $\widehat{\mathbf{W}}_{\text{SOUND}}$. In a parallel (asynchronous) process, the algorithm waits at least until the previous update of the SOUND filter has finished. In practice, we set an upper limit to the update rate, based on the available computational power. In the asynchronous process, we fetch the latest samples from the data buffer and baseline correct the fetched data. We then update the SOUND filter based on the latest data- and noise covariance estimates (see Eqs. S8, S13, and S15).

**Table S1.** A list of the tunable rtSOUND parameters, including their intended effects, chosen values in this study, and the rationale for each value.

| Parameter | Effect | Value | Rationale for chosen value |
|---|---|---|---|
| SOUND filter update interval | Retains the SOUND update rate within computational limits. | 100 ms (500 samples) | Well within computational limits, while still ensuring low-latency reaction to changes in noise distribution. |
| Length of data buffer | Balances between algorithm agility, stationarity assumptions, and computational speed (see Eq. S15). | 100 ms (500 samples) | Provides sufficient data for SOUND to reliably estimate the current noise distribution. |
| Pause after TMS pulse | Prevents SOUND from updating during a time window after a TMS pulse, to avoid contamination by pulse artifacts and TEP-related fluctuations. | 1 s | In 1 s, the TEP and particularly its artifacts are mostly over [7]. |
| Convergence boundary | Ensures the SOUND filter solution has converged adequately by limiting the maximum relative difference in channel-specific noise variances between consecutive iterations. | 0.01 | Suggested in [1, 2]. Cleaned data remained practically identical whether iteration was continued until full convergence or stopped at this 1% threshold. |
| Maximum number of iterations | Prevents indefinite iteration during the SOUND update procedure. | 10 | Suggested in [1, 2]. With typical numbers of EEG channels, the filter does not significantly change beyond 10 iterations. |
| Regularization level $\lambda_0$ | Tunes the trade-off between improving SNR and avoiding overcorrection (Eq. S6). | 0.1 | Suggested in [1]. This is a key parameter for tuning the SOUND performance. |
| Baseline update rate $\alpha$ | Prevents large, sudden baseline shifts, improving the robustness of SOUND (Eq. S19). | 0.0006 | This rate gives a half-attenuation time of approx. 200 ms. See Figure S2 for details. |
| Noise covariance update rate $\beta$ | Prevents abrupt changes between consecutive SOUND filters; balances between smoothness and responsiveness (Eq. S17). | 0.05 | Empirically found to produce a smooth signal. |

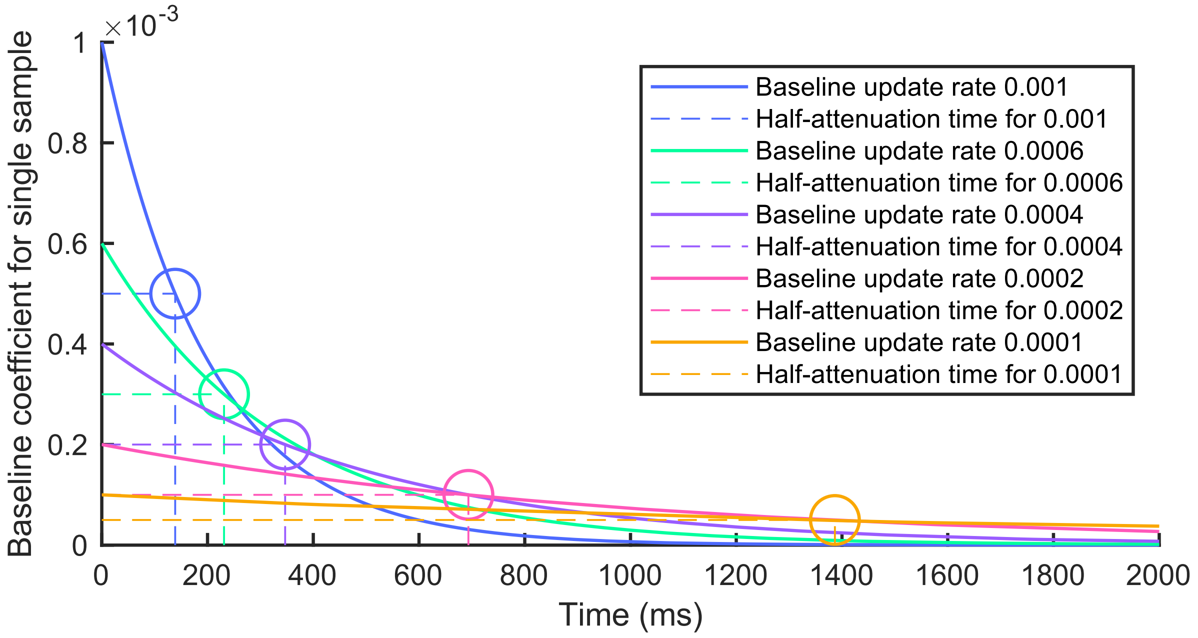


**Fig S2.** The real-time baseline correction was implemented as a geometric sum of the past samples. The baseline update rate determines how fast the baseline correction forgets past samples, *i.e.*, how fast the sample-specific baseline-correction coefficients decrease in time. The half-attenuation times denote the duration at which the effect (coefficient) of one sample on the baseline to decrease to half of its original value. The smaller the update rate, the longer each sample influences the baseline. The choice of the update rate is a trade-off between resistance against (small update rate) and fast recovery from (large update rate) baseline-disturbing artifacts.

### References

[1] Mutanen TP, Metsomaa J, Liljander S, Ilmoniemi RJ. Automatic and robust noise suppression in EEG and MEG: The SOUND algorithm. Neuroimage 2018;166:135-51.
[2] Mutanen TP, Metsomaa J, Makkonen M, Varone G, Marzetti L, Ilmoniemi RJ. Source-based artifact-rejection techniques for TMS–EEG. Journal of Neuroscience Methods 2022;382:109693.
[3] Weinstein D, Zhukov L, Johnson C. Lead-field bases for electroencephalography source imaging. Annals of biomedical engineering 2000;28:1059-65.
[4] Hämäläinen MS, Ilmoniemi RJ. Interpreting magnetic fields of the brain: minimum norm estimates. Medical & biological engineering & computing 1994;32:35-42.
[5] Allegrini P, Menicucci D, Bedini R, Fronzoni L, Gemignani A, Grigolini P, et al. Spontaneous brain activity as a source of ideal 1/f noise. Physical Review E 2009;80(6):061914.
[6] Metsomaa J, Song Y, Mutanen TP, Gordon PC, Ziemann U, Zrenner C, et al. Adapted Beamforming: A Robust and Flexible Approach for Removing Various Types of Artifacts from TMS–EEG Data. Brain Topography 2024:1-25.
[7] Hernandez-Pavon JC, Veniero D, Bergmann TO, Belardinelli P, Bortoletto M, Casarotto S, et al. TMS combined with EEG: Recommendations and open issues for data collection and analysis. Brain Stimulation 2023.
